# Complexome profiling of the Chlamydomonas *psb28* mutant reveals THYLAKOID ENRICHED FRACTION 5 as an early photosystem II assembly factor

**DOI:** 10.1101/2024.06.24.600430

**Authors:** Julia Lang, Katharina König, Benedikt Venn, Benjamin Spaniol, Lara Spaniol, Frederik Sommer, Matthieu Mustas, Stefan Geimer, Torben Fürtges, Pawel Brzezowski, Jure Zabret, Francis-André Wollman, Mark Nowacyzk, David Scheuring, Till Rudack, Timo Mühlhaus, Yves Choquet, Michael Schroda

**Affiliations:** Molekulare Biotechnologie & Systembiologie, RPTU Kaiserslautern-Landau, Paul-Ehrlich Straße 23, D-67663 Kaiserslautern, Germany; Computational Systems Biology, RPTU Kaiserslautern-Landau, Erwin-Schrödinger-Straße 56, D-67663 Kaiserslautern, Germany; Biologie du Chloroplaste et Perception de la Lumière chez les Microalgues, Institut de Biologie Physico-Chimique, UMR CNRS/UPMC 7141, Paris, France; Zellbiologie/Elektronenmikroskopie, Universität Bayreuth, 95440 Bayreuth, Germany; Structural Bioinformatics, Regensburg Center for Biochemistry, University of Regensburg, Universitätsstr. 31, D-93053 Regensburg, Germany; Humboldt-Universität zu Berlin, Lebenswissenschaftliche Fakultät, Institut für Biologie, AG Pflanzenphysiologie, 10115 Berlin, Germany; Department of Plant Biochemistry, Faculty of Biology and Biotechnology, Ruhr University Bochum, Bochum, Germany; Plant Pathology, RPTU Kaiserslautern-Landau, Paul-Ehrlich Straße 22, D-67663 Kaiserslautern, Germany

## Abstract

Several auxiliary factors are required for the assembly of photosystem (PS) II, one of which is Psb28. While the absence of Psb28 in cyanobacteria has little effect on PSII assembly, we show here that the Chlamydomonas *psb28*-null mutant is severely impaired in PSII assembly, showing drastically reduced PSII supercomplexes, dimers and monomers, while overaccumulating RCII, CP43_mod_ and D1_mod_. The mutant had less PSI and more Cyt*b_6_f* and showed fewer thylakoid stacks and distorted chloroplast morphology. Complexome profiling of the *psb28* mutant revealed that TEF5, the homolog of Arabidopsis PSB33/LIL8, co-migrated particularly with RCII. TEF5 also interacted with PSI. A Chlamydomonas *tef5* null mutant is also severely impaired in PSII assembly and overaccumulates RCII and CP43_mod_. RC47 was not detectable in the light-grown *tef5* mutant. Our data suggest a possible role for TEF5 in facilitating the assembly of CP47_mod_ into RCII. Both the *psb28* and *tef5* mutants exhibited decreased synthesis of CP47 and PsbH, suggesting negative feedback regulation possibly exerted by the accumulation of RCII and/or CP43_mod_ in both mutants. The strong effects of missing auxiliary factors on PSII assembly in Chlamydomonas suggest a more effective protein quality control system in this alga than in land plants and cyanobacteria.

**One-sentence summary:** The Chlamydomonas psb28 mutant is severely impaired in PSII assembly which via complexome profiling allowed identifying TEF5 as a novel PSII assembly factor that likely facilitates CP47 assembly.

The author responsible for distribution of materials integral to the findings presented in this article in accordance with the policy described in the Instructions for Authors (https://academic.oup.com/plcell/pages/General-Instructions) is: Michael Schroda.

## Introduction

Photosystem II (PSII) is a light-driven water-plastoquinone oxidoreductase in the thylakoid membranes of cyanobacteria and chloroplasts. Structural analyses of the PSII core complex from spinach and pea revealed four large intrinsic subunits, D1 (PsbA), D2 (PsbD), CP43 (PsbC) and CP47 (PsbB) as well as twelve small membrane-spanning subunits PsbE, PsbF, PsbH, PsbI, PsbJ, PsbK, PsbL, PsbM, PsbTc, PsbW, PsbX, and PsbZ. Moreover, there were four extrinsic subunits on the luminal side, including oxygen-evolving complex proteins (PsbO, PsbP, PsbQ) and PsbTn (Wei et al., 2016; Su et al., 2017). Structural analyses of PSII from *Chlamydomonas reinhardtii* (Chlamydomonas) revealed the same subunits as found in the PSII core from land plants but two more peripheral subunits were detected, Psb30 and PsbR, while PsbTn was absent. Moreover, two new densities referred to as unidentified stromal protein (USP) and small luminal protein (SLP) were detected (Sheng et al., 2019; Sheng et al., 2021). PSII core monomers assemble into dimers, to which peripheral antenna bind on both sides to form PSII supercomplexes. In land plants, a PSII dimer binds two of the monomeric minor antenna CP24 (LHCB6), CP26 (LHCB5) and CP29 (LHCB4) as well as up to four major LHCII heterotrimers (Caffarri et al., 2009; Kouril et al., 2011; Su et al., 2017). In Chlamydomonas, which lacks CP24, a PSII dimer binds two each of the CP26 and CP29 monomers as well as up to six large LHCII heterotrimers (Tokutsu et al., 2012; Sheng et al., 2019; Sheng et al., 2021).

Based largely on the seminal work on cyanobacterial PSII, the steps leading to the formation of PSII core complexes from assembly modules have been revealed (Nickelsen and Rengstl, 2013; Lu, 2016; Plochinger et al., 2016; Komenda et al., 2024): PSII assembly starts with the synthesis of the α- and β-subunits (PsbE and PsbF) of Cyt*b_559_*, which accumulates in the membrane and interacts with newly made D2 to form the D2_mod_ (Morais et al., 1998; Muller and Eichacker, 1999; Komenda et al., 2004). In parallel, the newly synthesized D1 precursor interacts with PsbI that has already been produced. PsbI-D1 (D1_mod_) is then combined with the D2-Cyt*b_559_* complex to form the reaction center (RCII) (Dobakova et al., 2007; Zhao et al., 2023). This is followed by proteolytic processing of the D1 precursor at its C-terminus (Anbudurai et al., 1994). With the low molecular mass subunits PsbH, PsbL, PsbM, PsbR, and PsbTc, CP47 forms the CP47_mod_ that combines with RCII to form the RC47 intermediate, which also contains PsbX and PsbY (Rokka et al., 2005; Boehm et al., 2012). CP43 interacts with the small subunits PsbK, PsbZ, and Psb30 and forms the CP43 module (CP43_mod_), which finally combines with RC47 to form PSII monomers (Sugimoto and Takahashi, 2003; Rokka et al., 2005; Boehm et al., 2011). During photoactivation in chloroplasts, the Mn_4_CaO_5_ cluster is attached to the luminal side of the PSII monomers, followed by the proteins PsbO, PsbP and PSBQ (Bricker et al., 2012). After dimerization and attachment of LHCII the assembly is complete and the supercomplex is transferred from stroma-exposed membranes to grana stacks (Tokutsu et al., 2012; van Bezouwen et al., 2017).

The assembly of PSII is facilitated by auxiliary factors that temporarily bind to discrete assembly intermediates, but they are not constituents of the final complex. Many, but not all of these auxiliary factors are conserved between cyanobacteria and chloroplasts (Nixon et al., 2010; Nickelsen and Rengstl, 2013; Lu, 2016; Komenda et al., 2024). For example, auxiliary factors including HCF136 (Ycf48 in *Syncechocystis*), PsbN, PAM68, PSB28, HCF244 (Ycf39 in *Synechocystis*) are conserved between chloroplasts and cyanobacteria, while factors such as Psb34 and Psb35 exist only in cyanobacteria and factors such as LPA2 exist only in chloroplasts. The conserved assembly factor Psb28 (Psb28-1 in *Synechocystis*) is peripherally associated at the cytoplasmic side mainly with RC47 and less with PSII monomers (Kashino et al., 2002; Dobakova et al., 2009; Sakata et al., 2013). Since Psb28 interacts with RC47 and PSII monomers only transiently, PSII complexes with Psb28 can be enriched in cyanobacterial mutants that accumulate PSII assembly intermediates, such as deletion mutants of *psbC* (Boehm et al., 2012), *psbJ* (Nowaczyk et al., 2012; Zabret et al., 2021) or *psbV* (Xiao et al., 2021). PSII assembled completely in the *Synechocystis psb28-1* mutant and was photochemically fully active (Dobakova et al., 2009; Sakata et al., 2013; Beckova et al., 2017). Accordingly, *psb28-1* showed no growth phenotype at various light intensities at 30°C, but a growth defect was observed at 38°C and light intensities of 30 µmol photons m^-2^ s^-1^ or higher (Sakata et al., 2013). Moreover, the *psb28-1* mutant was more sensitive to fluctuating light (Beckova et al., 2017). The isolation of tagged Psb28 from the cyanobacterial *psbJ* and *psbV* mutants allowed determining the cryo-EM structures of Psb28 bound to RC47 and PSII monomers (Xiao et al., 2021; Zabret et al., 2021). The Psb28-RC47 complex contained PSII subunits D1, D2, CP47, PsbE, PsbF, PsbH, PsbI, PsbL, PsbM, PsbT, and PsbX, while the Psb28-PSII monomer complex also contained the CP43_mod_ bound in a premature conformation. Psb28 was found to associate with D1, D2, and CP47 directly at the cytosolic surface of PSII. Psb28 binding induces the formation of an extended β-hairpin structure that incorporates Psb28’s central antiparallel β-sheet, the C terminus of CP47 and the D-E loop of D1. Psb28 binding causes large structural changes at the D–E loop regions of D1 and D2 when compared with native PSII, which affects the environment of the Q_A/B_ binding sites and the non-haem iron, potentially changing the Q_A_/Q_A_^-^ redox potential to reduce singlet oxygen production and thus prevent photodamage (Xiao et al., 2021; Zabret et al., 2021).

The function of some PSII auxiliary factors is less clear. An example is PSB33/LIL8, which interacts with RC47 and larger PSII assembly states, but mainly with PSII monomers, and locates to stroma lamellae and grana margins in *Arabidopsis thaliana* (Arabidopsis) (Fristedt et al., 2015; Fristedt et al., 2017; Kato et al., 2017; Nilsson et al., 2020). Arabidopsis *psb33/lil8* mutants gave rise to an “emergent” PSII phenotype that was only observed during a suite of varying light treatments over five days (Cruz et al., 2016), possibly explaining the very different mutant phenotypes reported. The “emergent” PSII phenotype was attributed to the formation of a fraction of PSII centers defective in Q_A_^-^ re-oxidation, possibly related to damage to the PSII Q_B_ site, which correlates with a more oxidized PQ pool reported by Kato et al. (2017).

Complexome profiling (CP) is based on the analysis of membrane protein complexes in gel bands of blue-native (BN) gels by mass spectrometry and can reveal novel assembly factors based on their co-migration with assembly intermediates (Heide et al., 2012; Heide and Wittig, 2013). We have previously employed CP on the Chlamydomonas *lpa2* mutant which allowed us to identify putative novel factors involved in PSII assembly steps beyond RCII (Spaniol et al., 2022). We found PSB28 to co-migrate with RC47 and PSII monomers in the *lpa2* mutant but not in WT, suggesting a conserved role of PSB28 in PSII assembly. Since PSB28 was not studied yet in molecular detail in chloroplasts, we characterized the Chlamydomonas *psb28* mutant. Unexpectedly, we found that the *psb28* mutant was strongly impaired in accumulating PSII assemblies beyond RCII, very much in contrast to cyanobacterial *psb28* mutants. We used CP on *psb28* and we found TEF5, the homolog of Arabidopsis PSB22/LIL8, to co-migrate with early PSII assembly intermediates, particularly RCII. We characterized the Chlamydomonas *tef5* mutant and we found that it is strongly affected in the accumulation of PSII, with hardly detectable RC47. This suggests a role of TEF5 in facilitating assembly of the CP47_mod_ into RCII. Overall, our results suggest that the absence of PSII auxiliary factors has a much greater impact on PSII assembly in Chlamydomonas than in cyanobacteria or land plants.

## Results

### The Chlamydomonas *psb28* mutant accumulates less PSII and PSI subunits and shows impaired growth in high light and under photoautotrophic conditions

*Synechocystis sp. (*strain *PCC 6803)* is equipped with two functionally distinct Psb28 homologs, Psb28-1 and Psb28-2 (Dobakova et al., 2009; Sakata et al., 2013; Beckova et al., 2017), while only single PSB28 proteins exist in Arabidopsis and Chlamydomonas. As shown in Figure 1A, Chlamydomonas and Arabidopsis PSB28 proteins are more closely related to *Synechocystis* Psb28-1 (58/71% and 45/69% identical/similar amino acid residues, respectively) than to Psb28-2 (28/45% and 34/54% identical/similar amino acid residues, respectively) and harbor predicted chloroplast transit peptides. The structure of Chlamydomonas PSB28 predicted by AlphaFold is very similar to that determined for *Thermosynechococcus elongatus* bound to the PSII acceptor side (RMSD = 1.24 Å; TM score = 0.91; Figure 1B) (Zabret et al., 2021).

**Figure 1.**
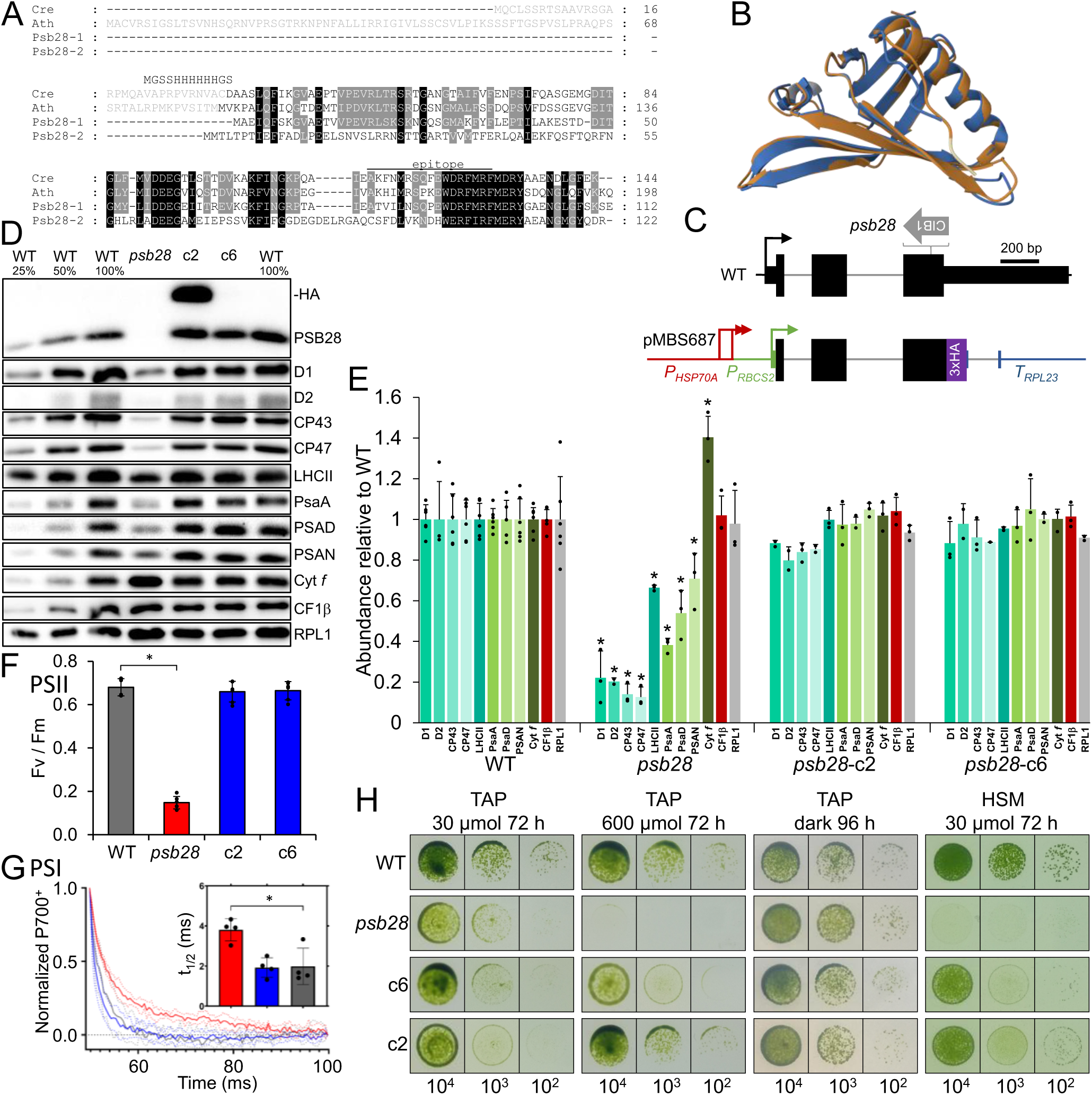
Phenotypes of the *psb28* mutant compared to WT and complemented lines. **(A)** Alignment of PSB28 amino acid sequences from *Chlamydomonas*, *Arabidopsis*, and *Synechocystis*. Residues highlighted in black and gray are conserved in four and three of the sequences, respectively. Predicted chloroplast transit peptides are shown in gray. The sequence with a hexahistidine tag replacing the transit peptide for production of recombinant *Chlamydomonas* PSB28 is shown. The peptide from *Chlamydomonas* PSB28 used for antibody production is indicated by a horizontal line. Ath – *Arabidopsis thaliana* (AT4G28660), Cre – *Chlamydomonas reinhardtii* (Cre10.g440450), Psb28-1 – *Synechocystis sp.* PCC 6803 variant 1 (Sll1398), Psb28-2 – *Synechocystis sp.* variant 2 (Slr1739). **(B)** Pairwise structure alignment of Psb28 from *T. elongatus* in its conformation when binding to the PSII acceptor side (7NHQ) (gold) and the AlphaFold structure of *Chlamydomonas* PSB28 lacking the chloroplast transit peptide (blue). **(C)** Structure of the *Chlamydomonas PSB28* gene, insertion site of the CIB1 cassette in the *psb28* mutant, and construct for complementation. Protein coding regions are drawn as black and purple boxes, untranslated regions as bars, and introns and promoter regions as thin lines. Arrows indicate transcriptional start sites. **(D)** Immunoblot analysis of the accumulation of PSB28 and of subunits of the major thylakoid membrane protein complexes. c2 and c6 are lines complemented with the construct shown in (C). PSII – D1, D2, CP43, CP47, LHCII; PSI – PsaA, PSAD, PSAN; Cyt *b_6_f* complex – Cyt *f*; ATP synthase – CF1β. Ribosomal protein RPL1 served as loading control. 10 µg of whole-cell proteins (100%) were analysed. **(E)** Quantification of the immunoblot analysis shown in (D). Values are means from three independent experiments normalized first by the median of all signals obtained with a particular antiserum in the same experiment, and then by the mean signal of the WT. Error bars represent standard deviation. Asterisks indicate significant differences with respect to the WT (two-tailed, unpaired *t*-test with Bonferroni-Holm correction, *P* < 0.05). The absence of an asterisk means that there were no significant differences. **(F)** F_v_/F_m_ values of the *psb28* mutant versus WT and complemented lines. Shown are averages from six independent experiments. Error bars represent standard deviation. The asterisk indicates significant differences between WT and *psb28* mutant/complemented lines (two-tailed, unpaired *t*-test with Bonferroni-Holm correction, *P* < 0.001). **(G)** PSI reduction kinetics of WT, *psb28* mutant and a complemented line. Shown are averages from four independent experiments fitted with single exponential functions. Standard deviations are shown as dotted lines. The asterisk indicates significant differences between WT and *psb28* mutant (one-way ANOVA, *P* < 0.01). **(H)** Analysis of the growth of 10^4^ – 10^2^ spotted cells under the conditions indicated.

The Chlamydomonas *psb28* mutant (Li et al., 2016) contains the CIB1 mutagenesis cassette within the third exon of the *PSB28* gene (Figure 1C). While we were able to amplify *PSB28* sequences flanking the cassette on the 5’ side by PCR, no PCR product was obtained on the 3’ side, presumably because flanking sequences were deleted, or a large piece of junk DNA had integrated between *PSB28* sequences and the CIB1 cassette (Supplemental Figure S1A, B). An antibody raised against a peptide from the C-terminal part of the *Chlamydomonas* PSB28 protein (Figure 1A) specifically detected a protein band at the expected molecular mass of ∼12.5 kDa in the wild type (WT), which was absent in the *psb28* CLiP mutant (Figure 1D; Supplemental Figure S1C). In the *psb28* mutant, PSII core subunits accumulated at most to 22% and LHCIIs to ∼66% of WT levels (Figures 1D, E). PSI core subunits accumulated to between 38% and 71% of WT levels. While the abundance of ATP synthase subunit CF1β was unaltered between mutant and WT, Cyt *f* of the Cyt *b_6_f* complex was 1.4-fold more abundant in the mutant compared with the WT. We amplified the genomic *PSB28* coding sequence, fused it with a sequence encoding a C-terminal 3xHA tag and placed it under control of the constitutive *HSP70A-RBCS2* promoter and the *RPL23* terminator using Modular Cloning (Figure 1C) (Schroda et al., 2000; Crozet et al., 2018). We combined the *PSB28* transcription unit with the *aadA* cassette (Meslet-Cladiere and Vallon, 2011) and transformed it into the *psb28* mutant. Seven picked spectinomycin resistant transformants that showed a greener appearance than the *psb28* mutant accumulated D1 roughly at WT levels and in all but two the HA tag was detected (Supplemental Figure S2; Figure 1D). Further analysis with the PSB28 peptide antibody of two transformants with (*psb28*-c2) and without (*psb28*-c6) detectable HA signal revealed that in both lines PSB28 accumulated to the WT level with the molecular mass corresponding to the WT protein (Figure 1D). In line *psb28*-c2, PSB28 with 3xHA tag accumulated additionally. These findings point to a processing of the 3xHA tag and to a controlled accumulation of the processed WT protein. The reduced accumulation of photosystem core subunits and LHCII as well as the increased accumulation of Cyt *f* were fully reversed in both complemented lines (Figure 1D, E). In accordance with the reduced accumulation of PSII core subunits, PSII maximum quantum yield, as indicated by the Fv/Fmax value, was strongly reduced in the *psb28* mutant (0.15) versus WT and complemented lines (0.66-0.68) (Figure 1F). The half-life of P700^+^ reduction was about twice as high in the *psb28* mutant compared to WT and a complemented line, indicating reduced electron flow through PSI in the mutant (Figure 1G). While the *psb28* mutant could grow under heterotrophic conditions and under mixotrophic conditions in low light (30 µmol photons m^-2^ s^-1^), it failed to grow under mixotrophic conditions in high light (600 µmol photons m^-2^ s^-1^) and under photoautotrophic conditions in low light (Figure 1H).

A *Synechocystis psb28-1* mutant was reported to accumulate magnesium protoporphyrin IX monomethyl ester and to contain a decreased level of protochlorophyllide, indicating inhibition of chlorophyll biosynthesis at the cyclization step and suggesting a role of Psb28-1 in regulating chlorophyll biosynthesis (Dobakova et al., 2009). Later work indicated that this phenotype was due to a defect in the strain background used for mutant construction (Beckova et al., 2017). To investigate whether PSB28 might regulate a specific step in chlorophyll (Chl) biosynthesis in Chlamydomonas, we measured the content of Chl a and b, of several Chl precursors, and of Chl breakdown product pheophorbide by HPLC in the WT and the *psb28* mutant grown in low light or in the dark for 65 h. All analyzed pigments accumulated to lower levels in the mutant when compared with the WT under both growth conditions, except for Chl a and Chl b, which accumulated to similarly low levels in the WT and the mutant grown in the dark (Supplemental Figure S3). This was observed also in spot tests of dark-grown cells (Figure 1H). Overall, the reduced levels of Chl and of all its precursors in the mutant versus the WT point to an overall reduced Chl synthesis in the mutant rather than to a specific block at a particular synthesis step.

### PSB28 is localized to the chloroplast, where its absence results in severe defects of chloroplast morphology and thylakoid ultrastructure

To investigate whether the absence of PSB28 affected cell morphology and thylakoid ultrastructure, we used light and transmission electron microscopy (TEM), respectively. Light microscopy revealed an abnormal chloroplast in the *psb28* mutant, with green areas restricted to the region around the pyrenoid and the distal part of the chloroplast lobes (Figure 2A). This phenotype was restored in the complemented lines. TEM revealed that the thylakoid membranes in the mutant are loosely arranged with stacks occurring only occasionally (Figure 2B). Using a mouse antibody against the HA tag and a rabbit antibody against D1, we determined the intracellular localization of PSB28-3xHA in the complemented line *psb28*-c2 by immunofluorescence. As shown in Figure 2C, PSB28-3xHA was detected in the cup-shaped chloroplast and co-localized with D1 in most areas, but there were also areas where PSB28-3xHA was present but not D1, particularly around the pyrenoid. Notice that only the fraction of PSB28 was detected which still contained a C-terminal 3xHA tag.

**Figure 2.**
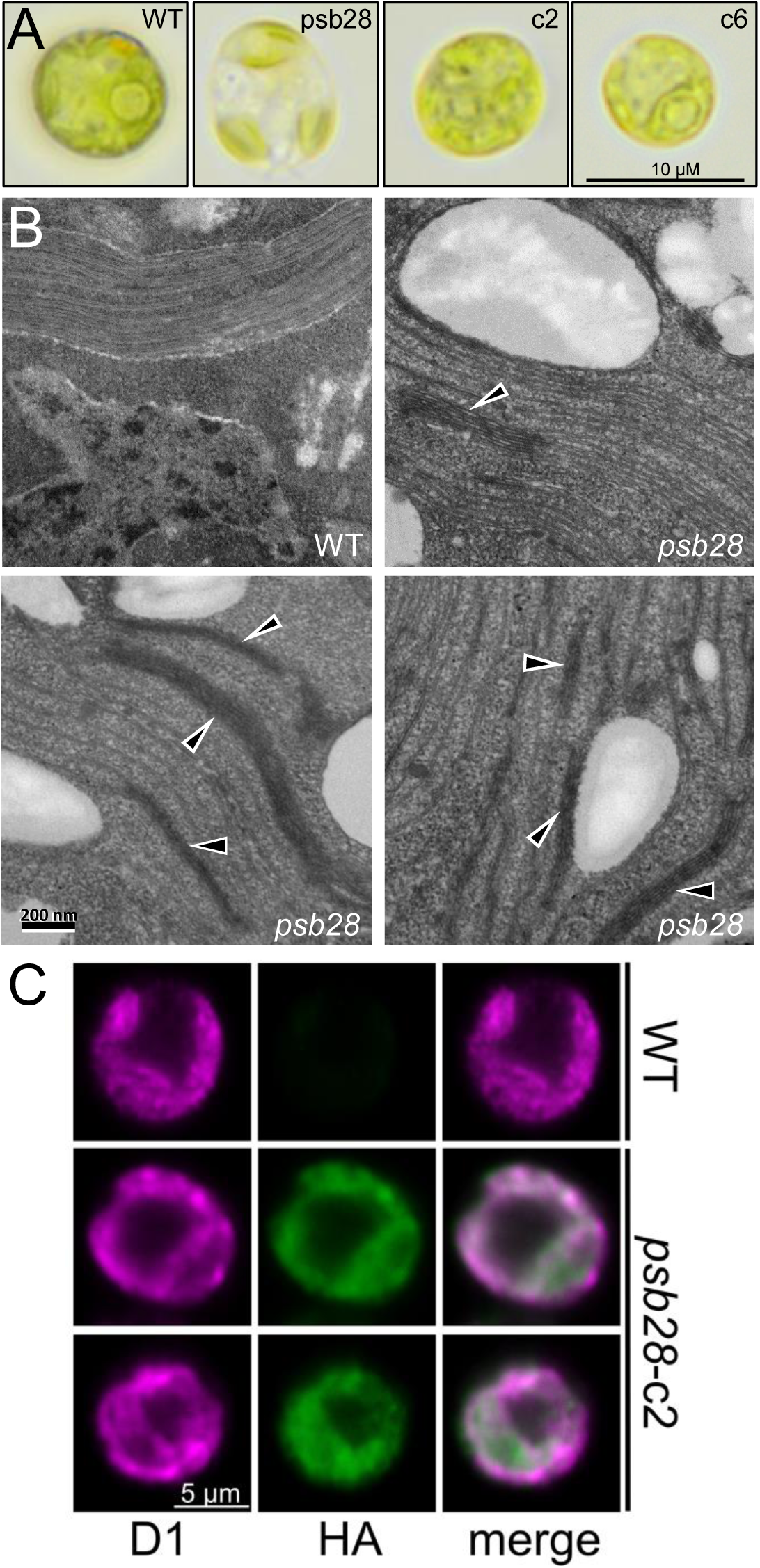
Light and electron microscopy of the *psb28* mutant and localization of PSB28 by immunofluorescence. **(A)** Light microscopy images of WT, *psb28* mutant, and complemented lines grown under mixotrophic conditions in low light (30 µmol photons m^-2^ s^-1^). **(B)** Electron microscopy images of WT and *psb28* mutant grown under mixotrophic conditions in low light. Black triangles indicate the rarely occurring thylakoid membrane stacks in the mutant. **(C)** Immunofluorescence localization of the D1 protein (magenta) and HA-tagged PSB28 (green) in a WT cell and two complemented *psb28* mutant cells (*psb28*-c2).

### The synthesis of subunits of both photosystems is impaired in the *psb28* mutant

To investigate effects of the lack of PSB28 on the synthesis and stability of newly made chloroplast-encoded photosynthetic proteins, we performed pulse-chase analyses with ^14^C-acetate on the WT, the *psb28* mutant, and complemented lines *psb28*-c2 and *psb28*-c6. Based on the ^14^C-labeling of proteins within the 7-min ^14^C-acetate pulse, the synthesis of PsaB, CP47, and PsbH was severely impaired in the *psb28* mutant (Figure 3). Synthesis and stability of D1, D2, and CP43 were reduced in the mutant, whereas synthesis and stability of RbcL and of subunits of the Cyt *b_6_f* complex and the ATP synthase were not affected when compared with the WT. These defects in the *psb28* mutant were fully restored in both complemented lines.

**Figure 3.**
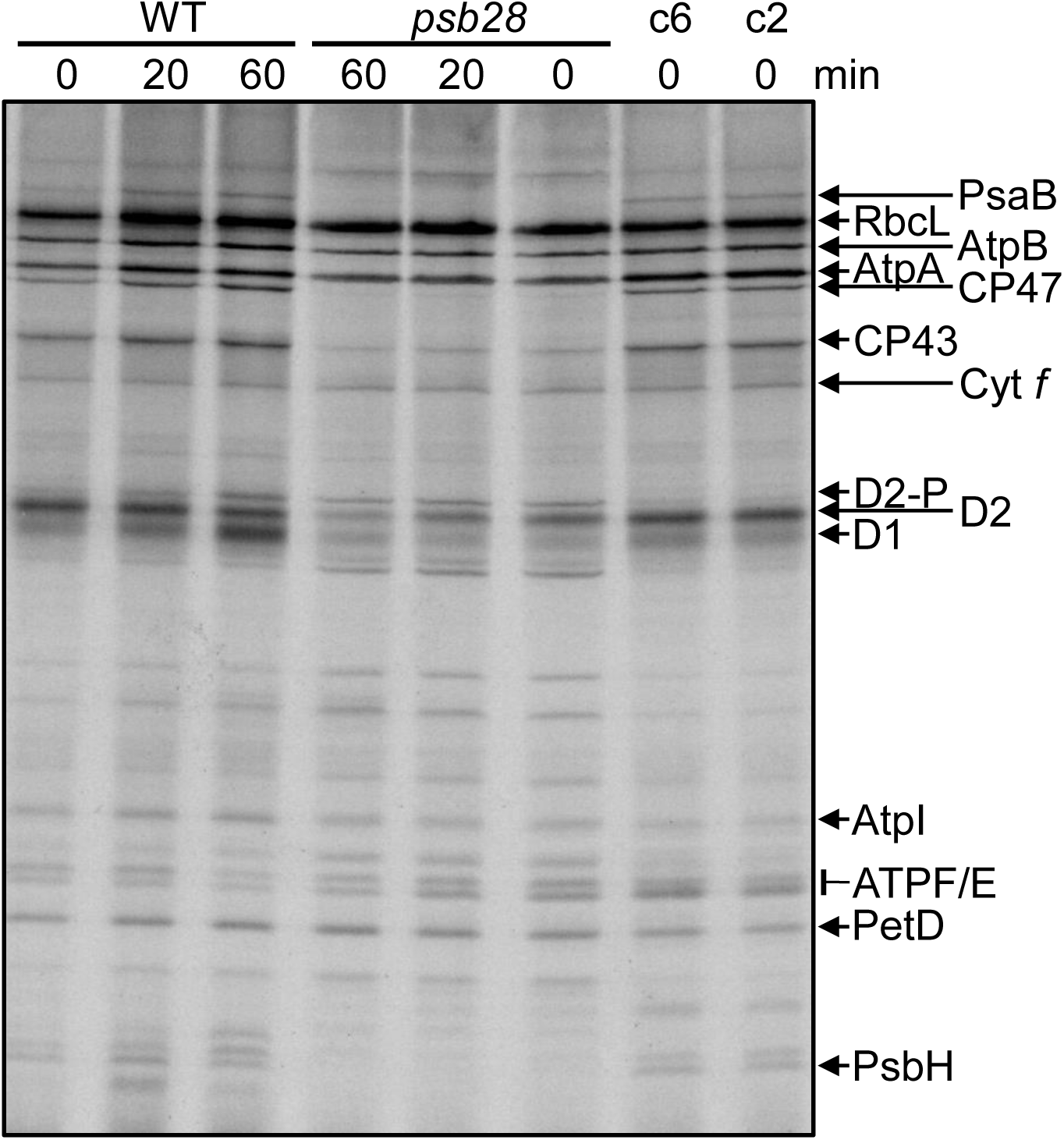
Analysis of synthesis and stability of thylakoid membrane proteins in the *psb28* mutant by pulse-chase labeling. WT, *psb28* mutant and complemented lines c2 and c6 were labelled with ^14^C-acetate in low light (20 µmol photons m^-2^ s^-1^) for 7 min in the presence of cytosolic translation inhibitor cycloheximide (0) and chased with unlabelled acetate for 20 and 60 min. Proteins were separated on a 12-18% SDS-urea gel and visualized by autoradiography. The assignment of the protein bands is based on mutant analyses (de Vitry et al., 1989; Girard-Bascou et al., 1992; Minai et al., 2006).

### PSII assembly states beyond RCII are severely reduced in the *psb28* mutant

To assess how the lack of PSB28 affects the PSII assembly states, we analyzed whole-cell protein extracts from the low light-grown WT, the *psb28* mutant, and the complemented lines by BN-PAGE and immunoblotting. While PSII supercomplexes, dimers, and monomers were detected with similar intensities in the WT and the complemented lines with antibodies against D1 and CP43, only a very faint signal for PSII monomers was detected in the *psb28* mutant with the D1 antibody (Figure 4A). However, we could detect RCII and CP43_mod_ in the mutant, which were not detectable in the WT and the complemented lines.

**Figure 4.**
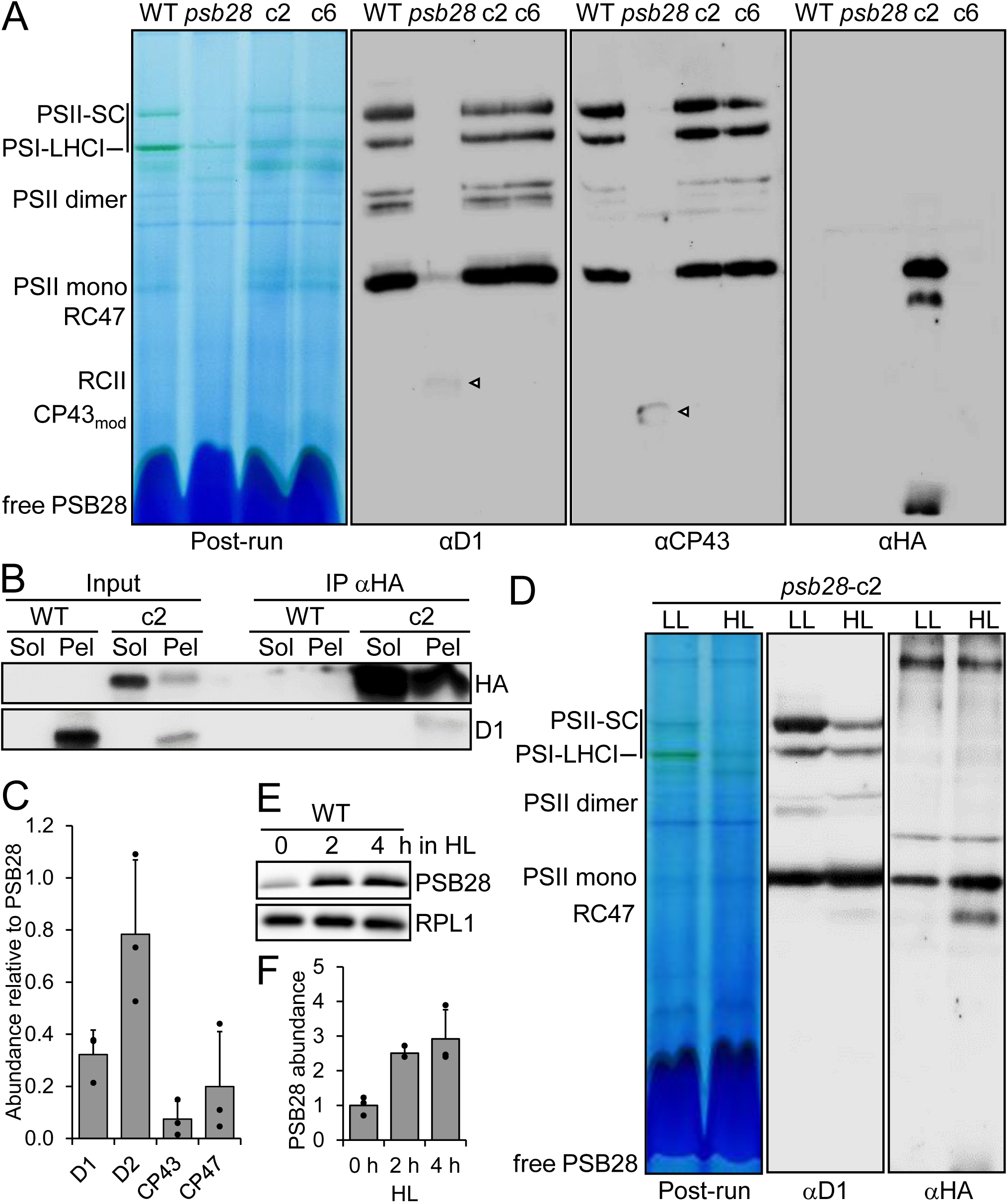
Analysis of protein complexes in the *psb28* mutant and of PSB28 interaction partners. **(A)** BN-PAGE analysis of proteins from cells grown in low light (30 µmol photons m^-2^ s^-1^). 50 µg of whole-cell proteins from WT, *psb28* mutant, and complemented lines *psb28*-c2 and *psb28*-c6 were solubilized with 1% β-DDM and separated on a 4-15% BN gel. Shown is a picture of the gel after the run and an immunoblot detected with antibodies against D1, CP43, and the HA epitope. Arrowheads point to faint bands likely representing RC47 and CP43_mod_ in the *psb28* mutant. SC – supercomplexes. **(B)** Immunoprecipitation of PSB28. Cells from complemented line *psb28*-c2 were fractionated via freeze-thaw cycles and centrifugation. HA-tagged PSB28 was then immunoprecipitated (IP) from soluble (Sol) and membrane-enriched (Pel) fractions with an HA antibody. 1% of the input and 10% of the precipitate were analysed by SDS-PAGE and immunoblotting using antibodies against the HA epitope and D1. **(C)** Mass spectrometry-based quantification of proteins co-precipitated from solubilized membrane fractions with HA-tagged PSB28. IBAQ values for each PSII core subunit were normalized by the IBAQ value for PSB28. Shown are mean values from three independent experiments. Error bars represent standard deviation. **(D)** BN-PAGE analysis of proteins from cells grown in low light (LL, 30 µmol photons m^-2^ s^-1^) and then exposed to high light (HL, 1200 µmol photons m^-2^ s^-1^) for 4 h. Whole-cell proteins from complemented line *psb28*-c2 were solubilized with 1% β-DDM and separated on a 4-15% BN gel. Shown is a picture of the gel after the run and an immunoblot detected with antibodies against D1 and the HA epitope. **(E)** Analysis of PSB28 accumulation in high light (HL). WT was exposed to 1200 µmol photons m^-2^ s^-1^ for 4 h and samples taken prior, 2 and 4 h after the treatment were analysed by immunoblotting using the peptide antibody against PSB28 and an antibody against RPL1 as loading control. **(F)** Quantification of the immunoblot analysis shown in (E). Values are means from three independent experiments. Normalization was done as described for Figure 1D.

We wondered whether the reduced accumulation of PSII and PSI core subunits and the reduced accumulation of PSII assembly states, beyond the RCII in the *psb28* mutant, were due to light-induced damage. To test this, we compared protein complexes in solubilized whole-cell extracts from the WT and the *psb28* mutant grown in low light and in the dark for 45 h. We observed an equally impaired accumulation of PSII and PSI complexes in the *psb28* mutant under both growth conditions (Supplemental Figure S4A). As determined by SDS-PAGE and immunoblotting, the accumulation of D1, CP43, and PsaA in the mutant was similarly affected in low light and in the dark (Supplemental Figure S4B). Nevertheless, the Fv/Fm value in the *psb28* mutant was slightly higher in dark-grown versus light-grown cells (0.2 vs 0.12, *P* = 0.014), while the opposite was observed for the WT (0.66 vs. 0.53, *P* = 0.001) (Supplemental Figure S4C). We conclude that the reduced accumulation of photosystems in the *psb28* mutant is not caused by damage inflicted by light.

### PSB28 interacts with complexes containing D2, D1, CP47 and CP43

Detection with the HA antibody revealed that PSB28-3xHA co-migrated with PSII monomers and, to a lesser extent, with RC47 in the *psb28*-c2 line (Figure 4A) (we did not use the antibody against PSB28 because it showed many cross-reactions with other proteins (Supplemental Figure 1C)). This suggests that excess PSB28-3xHA contributes to the pool of functional PSB28 in this line. A substantial fraction of free PSB28-3xHA was detected, as well. To rule out that PSB28 forms oligomers that co-migrate with PSII monomers and RC47 by chance, we analyzed migration properties of recombinantly produced PSB28 on BN gels. Recombinant PSB28 migrated entirely below the ∼25-kDa monomeric nucleotide exchange factor CGE1 (Willmund et al., 2007), indicating that *Chlamydomonas* PSB28 forms at most dimers but no higher oligomers (Supplemental Figure S5A). We also used recombinant PSB28 to estimate its abundance in the cell by quantitative immunoblotting and found that PSB28 constitutes 0.0034 ± 0.001% of the total protein content in the cell (Supplemental Figure S5B). Assuming ∼25 pg of total protein/cell, PSB28 would make up to 0.07 attomol/cell. Compared with an estimated 5.2 attomol PSII/cell, PSB28 would be ∼78-fold less abundant than PSII (Hammel et al., 2018; Hammel et al., 2020).

To verify the interaction of PSB28 with RC47/PSII monomers that was implied from their co-migration in BN-PAGE, we used the HA antibody to immunoprecipitate PSB28-3xHA from soluble and membrane-enriched fractions prepared from the complemented *psb28*-c2 line. Prior to immunoprecipitation, complexes were stabilized by *in-vivo* crosslinking with 0.37% formaldehyde. Immunoblot analyses showed that more HA-tagged PSB28 was immunoprecipitated from the soluble fraction than from the membrane fraction (Figure 4B). Moreover, D1 was detected only in PSB28 precipitates generated from membrane fractions. To identify and quantify all of the proteins interacting with PSB28, we analyzed the PSB28 immunopreciptates by LC-MS/MS (Supplemental Dataset S1). In precipitates from soluble fractions, PSB28 was the only protein detected in all three replicates. In precipitates from the membrane fractions, only D1, D2, CP43, and CP47 were detected in all three replicates, in addition to PSB28. Intensity-based absolute quantification (IBAQ) normalized to PSB28 revealed that D2 was the most abundant protein in the precipitate, followed by D1, CP47, and CP43 (Figure 4C).

### PSB28 abundance and its association with PSII increase in high light

*Synechocystis* PSB28-1 and 2 were found in PSII-PSI supercomplexes particularly under high-light intensities (Beckova et al., 2017). To test whether this is true also for Chlamydomonas PSB28, we exposed the complemented line *psb28*-c2 to low and high light intensities, solubilized whole-cell proteins, separated them by BN-PAGE, and detected D1 and HA-tagged PSB28 by immunoblotting (Figure 4D). Based on the D1 signals, more RC47 and PSII monomers accumulated at the expense of PSII supercomplexes in high versus low light. More PSB28-3xHA was associated with RC47 and PSII monomers in high versus low light but we did not observe an increased association of PSB28-3xHA with larger complexes. However, we found a ∼2.9-fold accumulation of PSB28 protein in WT exposed to 1200 µmol photons m^-2^ s^-1^ for 4 h, pointing to a potential role of PSB28 in PSII repair (Figure 4E, F).

To investigate the susceptibility of PSII in the *psb28* mutant to high light and its capability to recover functional PSII, we exposed WT, *psb28* mutant, and the complemented lines to high light (1800 µmol photons m^-2^ s^-1^) in the presence of translation inhibitor chloramphenicol (CAP) for one hour and allowed the cells to recover in the presence and absence of CAP at low light (30 µmol photons m^-2^ s^-1^). All four lines recovered full initial PSII activity (and D1 protein levels) at similar rates within five hours in a protein synthesis-dependent manner (Supplemental Figure S6). We also monitored kinetics of PSII degradation and resynthesis in sulfur-depleted and sulfur-repleted cultures, respectively (Malnoe et al., 2014). Here, the *psb28* mutant lost PSII activity upon sulfur depletion faster than the WT and the complemented lines but recovered initial PSII activity (and D1 levels) with similar rates as the other lines (Supplemental Figure S7). In summary, the very low levels of PSII in *psb28* are susceptible to photoinhibition and degradation upon sulfur deprivation but can be fully recovered at WT rates.

### Psb28-1 from *Synechocystis* partially complements the Chlamydomonas *psb28* mutant

Given the similarity between PSB28 from Chlamydomonas and Psb28-1 from *Synechocystis* (Figure 1A, B) we attempted to complement the Chlamydomonas *psb28* mutant with *Synechocystis* Psb28-1. For this, we synthesized the coding sequence for Psb28-1 with optimal Chlamydomonas codon usage and inserted *RBCS2* intron 1 into the coding sequence to enhance gene expression (Baier et al., 2018; Schroda, 2019). We then fused the *psb28-1* gene with sequences encoding the CDJ1 chloroplast transit peptide (Niemeyer et al., 2021) as well as a C-terminal 3xHA tag and placed it under control of the constitutive *HSP70A-RBCS2* promoter and the *RPL23* terminator using Modular Cloning (Supplemental Figure S8A). We combined the *psb28-1* transcription unit with the *aadA* cassette and transformed it into the *psb28* mutant. Twelve spectinomycin-resistant transformants (cs, <u>c</u>omplemented with *<u>S</u>ynechocystis* Psb28-1) were analyzed for the production of the recombinant protein by immunoblotting using an HA antibody, but specific signals could not be detected (not shown). We then monitored Fv/Fm values in liquid cultures and found that seven transformants had Fv/Fm values around or even below that of the *psb28* mutant but five had values that were significantly higher (P < 0.05) (Supplemental Figure S8B). The two transformants with the highest Fv/Fm values were cs9 and cs11 with values of 0.31 and 0.27, respectively, versus 0.2 for the *psb28* mutant (Figure 5A). Light microscopy revealed that transformants with Fv/Fm values below that of the *psb28* mutant showed the same defect in chloroplast morphology as the *psb28* mutant. However, transformants cs9 and cs11 had a WT chloroplast morphology (Figure 5B; Supplemental Figure S8C). Compared with the *psb28* mutant, both cs9 and cs11 showed improved growth under mixotrophic and photoautotrophic conditions in low light (30 µmol photons m^-2^ s^-1^) and were less sensitive to high light intensities (600 µmol photons m^-2^ s^-1^), but still fell far short of WT performance (Figure 5B). Immunoblot analyses revealed slightly higher levels of D1, D2, CP43, CP47, and PsaA in cs9 and cs11 than in the *psb28* mutant while levels of Cyt *f* remained high (Figure 5C). Accordingly, as revealed by BN-PAGE and immunoblotting, PSII monomers, dimers, and supercomplexes as well as PSI-LHCI were clearly more abundant in cs9 and cs11 than in the *psb28* mutant (Figure 5D).

**Figure 5.**
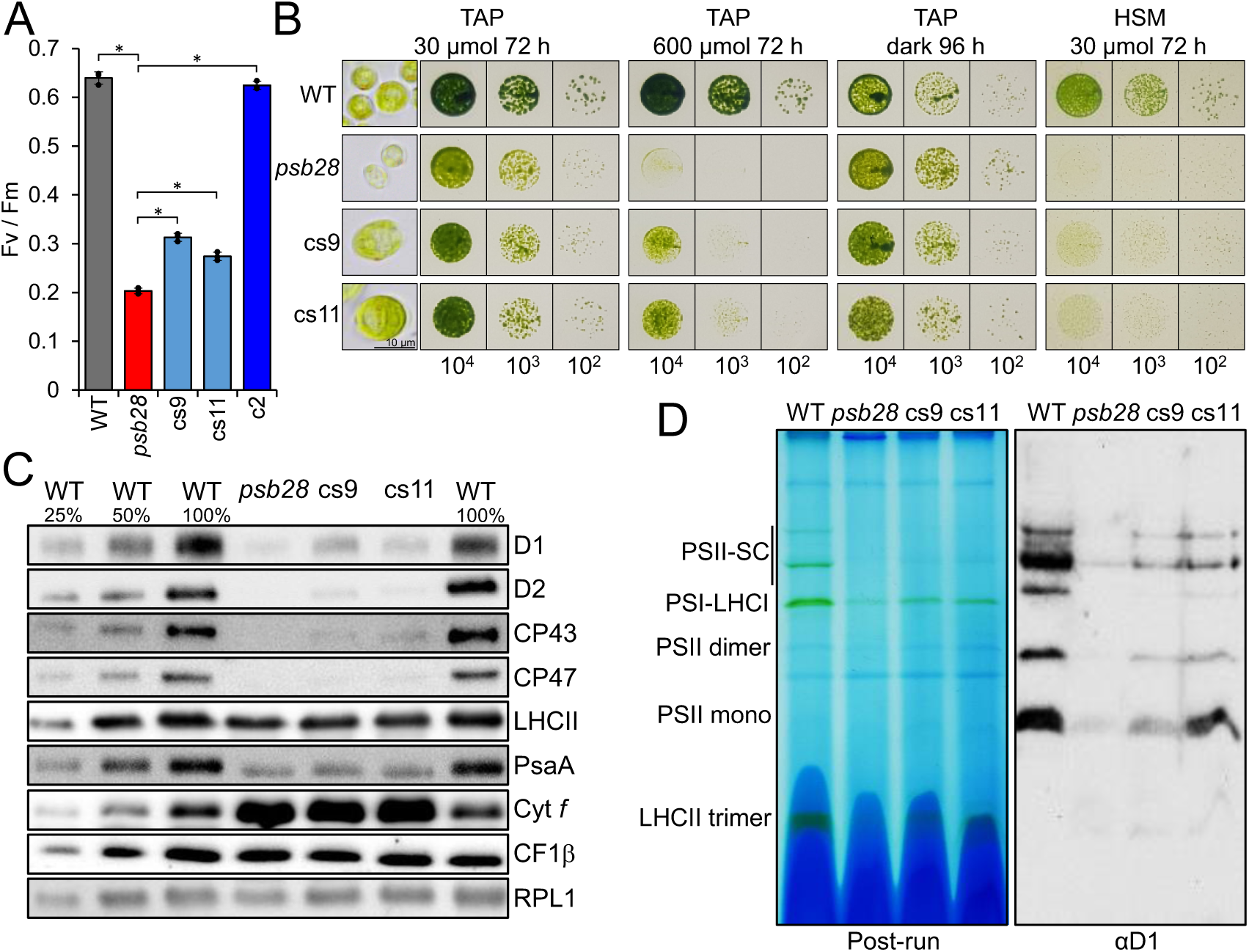
Complementation of the Chlamydomonas *psb28* mutant with *Synechocystis* Psb28-1. **(A)** F_v_/F_m_ values of the *psb28* mutant versus WT and lines complemented with Chlamydomonas PSB28 (c2) and *Synechocystis* Psb28-1 (cs9, cs11). Shown are averages from three independent experiments. Error bars represent standard deviation. Asterisks indicate significant differences with respect to the *psb28* mutant (two-tailed, unpaired *t*-test with Bonferroni-Holm correction, *P* < 0.001). **(B)** Light microscopy (left) and growth analysis of 10^4^ – 10^2^ spotted cells under the conditions indicated. **(C)** Immunoblot analysis of the accumulation of subunits of the major thylakoid membrane protein complexes. PSII – D1, D2, CP43, CP47, LHCII; PSI – PsaA; Cyt *b_6_f* complex – Cyt *f*; ATP synthase – CF1b. Ribosomal protein RPL1 served as loading control. 10 µg of whole-cell proteins (100%) were analysed. **(D)** BN-PAGE analysis. Cells of WT, *psb28* mutant, and complemented lines cs9 and cs11 were grown in low light (30 µmol photons m^-2^ s^-1^) and solubilized with 1% β-DDM. 60 µg of protein per lane were separated on a 4-15% BN gel. Shown is a picture of the gel after the run and an immunoblot detected with an antibody against D1.

In summary, *Synechocystis* Psb28-1 complements the Chlamydomonas *psb28* mutant, but with low efficiency. This could be due to its very low abundance, presumably caused by the instability of the protein, as we failed to detect the HA-tagged protein. Alternatively, as observed for Chlamydomonas PSB28, the HA tag could have been cleaved off and recombinant Psb28-1 accumulated in sufficient amounts but cannot fully complement the lack of the native PSB28.

### Complexome profiling of the *psb28* mutant reveals severe defects in PSII assembly beyond RCII

The accumulation of early PSII assembly intermediates in the *psb28* mutant prompted us to employ complexome profiling (CP) (Heide et al., 2012; Spaniol et al., 2022) to identify early PSII assembly factors by their co-migration with early PSII assembly intermediates. The analyses were performed on isolated thylakoid membranes from WT and *psb28* grown in low light (∼30 µmol photons m^-2^ s^-1^) in three biological replicates. Thylakoid membranes were solubilized with n-dodecyl α-D-maltoside (α-DDM) and protein complexes were separated on a 4-15% BN gel (Supplemental Figure S9). Each gel lane was cut into 36 slices and the resulting 216 slices were subjected to tryptic in-gel digestion and LC-MS/MS analysis. In total, 962 proteins were identified. Summed extracted ion chromatograms (XICs) of all peptides measured for a protein were used for protein quantification. To account for unequal loading, a normalization step was required. Thylakoid membranes from the *psb28* mutant lack most of PSII and part of LHCII and PSI (Figure 1D, E). Moreover, they are distorted (Figure 2A, B) and might behave differently from WT thylakoids during the extraction. Hence, normalization based on total ion intensities per lane, as done previously for CP on the *lpa2* mutant (Spaniol et al., 2022), appeared inappropriate. We therefore decided to normalize on the summed ion intensities of the eight identified ATPase subunits, as the abundance of the ATPase appeared unaffected when whole-cell proteins from *psb28* and WT were compared (Figure 1D, E). Ion intensity profiles for each protein along the BN gel run can be displayed from the interactive Excel table in Supplemental Dataset S2. The migration profiles of all identified proteins of WT and *psb28* mutant, clustered according to their migration behavior, are shown in Supplemental Dataset S3 as heat maps. The profiles for proteins belonging to the major thylakoid membrane complexes from WT and *psb28* are shown as heat maps in Figure 6A. Missing subunits, such as PsbI, did not give rise to detectable peptides because peptides are too small, too large, too hydrophobic or contain posttranslational modifications other than methionine oxidation or N-acetylation.

**Figure 6.**
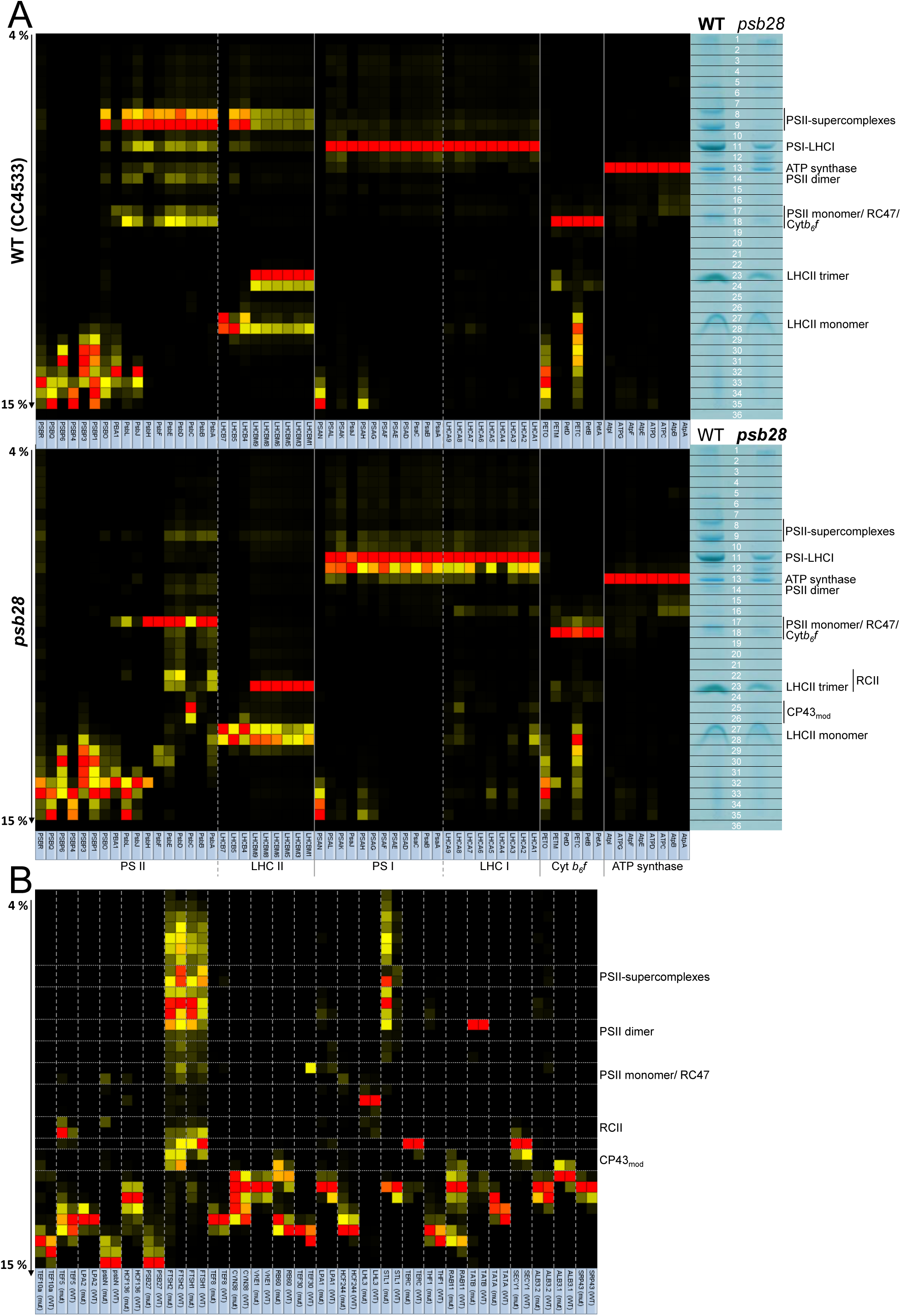
Complexome profiling on WT and *psb28* mutant. **(A)** Heat map showing the BN-PAGE migration profiles of subunits of the major thylakoid membrane protein complexes of WT (top panel) and *psb28* mutant (bottom panel). Values for each protein are derived from averaged peptide ion intensities from three biological replicates and are normalized to the gel slice with highest intensities. The BN-PAGE lane of one replicate from WT and *psb28* mutant is shown with the excised band corresponding to the heat map row. The underlying data and the migration profiles for each protein are accessible in Supplemental Dataset S2. **(B)** Heat map showing the BN-PAGE migration profiles of known and putatively new auxiliary factors involved in PSII biogenesis, repair, and the regulation of PSII complex dynamics in WT and *psb28* mutant (Supplemental Table S2).

Eight subunits of the ATP synthase and six subunits of the Cyt *b_6_f* complex were identified. The median abundance of the Cyt *b_6_f* complex was ∼1.87-fold higher in the *psb28* mutant compared to the WT (Table 1). Nevertheless, there were no differences in the migration patterns of ATPase and Cyt *b_6_f* complex subunits in WT and *psb28* mutant (Figure 6A). As reported previously, PETO did not interact stably with other subunits of the Cyt *b_6_f* complex (Takahashi et al., 2016) and a substantial fraction of the Rieske iron-sulfur protein migrated as unassembled protein (Spaniol et al., 2022).

**Table 1.** Ratio of subunit abundance between *psb28* mutant and WT. Values are based on the summed ion intensities in all gel bands of three biological replicates each of WT and mutant.

| ATP synthase |  | Cytb<sub>6</sub>f |  | Photosystem II |  | Photosystem I |  |
| --- | --- | --- | --- | --- | --- | --- | --- |
| Subunit | Ratio | Subunit | Ratio | Subunit | Ratio | Subunit | Ratio |
| AtpA | 1.03 | PetA | 1.97 | PsbA (D1) | 0.15 | PsaA | 1.03 |
| AtpB | 1.07 | PetB | 1.86 | PsbB (CP47) | 0.08 | PsaB | 0.80 |
| ATPC | 0.91 | PETC | 1.99 | PsbC (CP43) | 0.21 | PsaC | 0.70 |
| ATPD | 1.08 | PetD | 1.69 | PsbD (D2) | 0.21 | PSAD | 0.95 |
| AtpE | 0.86 | PETM | 1.88 | PsbE | 0.15 | PSAE | 0.57 |
| AtpF | 1.06 | PETO | 0.81 | PsbF | 0.10 | PSAF | 0.96 |
| ATPG | 1.03 | **Median** | **1.87** | PsbH | 0.005 | PSAG | 1.06 |
| AtpI | 0.82 |  |  | PsbJ | 0.42 | PSAH | 0.88 |
| **Median** | **1.03** |  |  | PsbL | 0.19 | PsaJ | 0.30 |
|  |  |  |  | PBA1 | 1.26 | PSAK | 0.75 |
|  |  |  |  | **Median** | **0.15** | PSAL | 0.90 |
|  |  |  |  | PSBO | 0.26 | PSAN | 0.50 |
|  |  |  |  | PSBP1 | 0.45 | **Median** | **0.84** |
|  |  |  |  | PSBP3 | 14.83 | LHCA1 | 0.81 |
|  |  |  |  | PSBP4 | 1.46 | LHCA2 | 0.60 |
|  |  |  |  | PSBP6 | 1.96 | LHCA3 | 0.89 |
|  |  |  |  | PSBQ | 0.35 | LHCA4 | 0.35 |
|  |  |  |  | PSBR | 0.27 | LHCA5 | 0.89 |
|  |  |  |  | **Median** | **0.45** | LHCA6 | 0.42 |
|  |  |  |  | LHCB4 | 0.61 | LHCA7 | 1.12 |
|  |  |  |  | LHCB5 | 1.24 | LHCA8 | 1.26 |
|  |  |  |  | LHCB7 | 5.13 | LHCA9 | 0.62 |
|  |  |  |  | LHCBM1 | 0.58 | **Median** | **0.81** |
|  |  |  |  | LHCBM3 | 0.47 |  |  |
|  |  |  |  | LHCBM5 | 0.55 |  |  |
|  |  |  |  | LHCBM6 | 0.46 |  |  |
|  |  |  |  | LHCBM8 | 0.44 |  |  |
|  |  |  |  | LHCBM9 | 0.43 |  |  |
|  |  |  |  | **Median** | **0.55** |  |  |

In contrast to the ATP synthase and the Cyt *b_6_f* complex, the composition of PSI differed between WT and mutant. While in WT only a single PSI-LHCI complex with eleven detected core subunits and nine LHCAs was observed, the mutant showed two prominent PSI-LHCI complexes that differed by the presence or absence of LHCA4 and LHCA6 (Figure 6A). As observed previously (Spaniol et al., 2022), some PSAH and all PSAN accumulated as unassembled subunits in both, *psb28* mutant and WT, presumably because they lost connection to the PSI core during sample preparation or electrophoresis. The median abundance of PSI core subunits and LHCI antennae was ∼16% and 19% lower in the mutant compared with the WT (Table 1).

The most dramatic change between *psb28* and WT was at the level of larger PSII complexes, with supercomplexes, dimers, and monomers/RC47 accumulating in the mutant only to 1%, 6%, and 27%, respectively, of WT levels, as judged from the median abundance of the core subunits in the complexes (Table 2; Figure 6A; Supplemental Figure S10). In contrast, D1 and D2 in RCII accumulated to more than 30-fold and CP43 in the CP43_mod_ to 10.5-fold higher levels in the mutant compared with the WT. D1 in D1_mod_ and PsbE in unassembled PsbE/F also accumulated 2.2- and 15.5-fold in the mutant, respectively. Overall, the median abundance of PSII core subunits in the *psb28* mutant was only ∼15% of that in the WT (Table 1). Even less CP47 and PsbH (∼8% and ∼0.5%, respectively, of WT levels) accumulated in the mutant, in line with their substantially lower synthesis rates (Figure 3). In contrast, the previously identified novel PSII-associated protein PBA1 (Spaniol et al., 2022) accumulated to 1.26-fold higher levels in the mutant compared to the WT, indicating that its abundance is not co-regulated with the canonical PSII core subunits. Except for PSBO, all other subunits involved in stabilizing/shielding the Mn_4_CaO_5_ cluster were found to migrate as unassembled subunits in both, *psb28* mutant and WT, presumably because they got detached from PSII during sample preparation or electrophoresis (Figure 6A). The median abundance of all subunits of the water-splitting complex reached ∼45% of WT levels (Table 1). Only PSBP3, 4 and 6 behaved differently and were 14.83-, 1.46- and 1.96-fold more abundant, respectively, in the mutant than in the WT. Hence, like PBA1, these three proteins are not co-regulated with the other PSII subunits. The mean abundance of LHCII proteins in the mutant reached only ∼55% of the values of the WT. (Table 1) and since hardly any larger PSII complexes were made in the mutant, it must contain a large pool of unassembled LHCII trimers and monomers. Only LHCB5 (CP26) and the recently identified LHCB7 protein (Klimmek et al., 2006) accumulated ∼1.24- and 5.13-fold, respectively, in the mutant when compared with the WT. In contrast to LHCB4 (CP29) and LHCB5, LHCB7 accumulated in the WT only in the unassembled form (Figure 6A), as observed previously (Spaniol et al., 2022).

**Table 2.** Ratio of subunit abundance in various PSII assembly states between *psb28* mutant and WT. Values are based on summed ion intensities in the bands indicated. SC – supercomplexes. RCII – reaction centers. ND, nd – not detected in WT, mutant (ion intensity < 0.05% of total intensity in respective strain).

|  | **SC** | **Dimers** | **Monomers/**<br>**RC47** | **RCII** | **CP43<sub>mod</sub>** | **D1<sub>mod</sub>/**<br>**PsbE/F** |
| --- | --- | --- | --- | --- | --- | --- |
| PsbA (D1) | 0.01 | 0.09 | 0.32 | 32.2 | ND | 2.2 |
| PsbB (CP47) | 0.01 | 0.07 | 0.28 | ND | ND | nd/ND |
| PsbC (CP43) | 0.02 | 0.09 | 0.24 | ND | 10.5 | ND |
| PsbD (D2) | 0.03 | 0.12 | 0.4 | 77.1 | ND | ND |
| PsbE | 0.02 | 0.05 | 0.26 | ND | ND | 15.5 |
| PsbF | nd | nd | 0.5 | nd/ND | nd/ND | ND |
| PsbH | nd | nd | 0.11 | nd/ND | nd/ND | nd/ND |
| PsbJ | 0.01 | 0.15 | 0.14 | nd/ND | nd/ND | nd/ND |
| PsbL | nd | 0.01 | 0.09 | ND | nd/ND | nd/ND |
| PSBO | 0.01 | 0.02 | nd | nd | nd | 0.6 |
| PBA1 | nd | nd/ND | 0.83 | nd/ND | nd/ND | 1.3 |
| **Median** | **0.01** | **0.06** | **0.27** |  |  |  |
| Gel bands | 7-11 | 13-15 | 17/18 | 22/23 | 25/26 | 28-30 |

When comparing mass spectrometry data of isolated thylakoids (Table 1) with whole cell immunoblot data (Figure 1D, E), we detected relatively more Cyt *b_6_f* and PSI in the *psb28* mutant than in the WT, but less PSII and LHCII. While we cannot exclude the possibility that the growth conditions used for the two data sets varied slightly (e.g. culture volume and perceived light), these differences could also be due to an unequal extractability of thylakoid membranes caused by the differences in thylakoid structure and composition between the mutant and the WT (Figure 2A, B).

### The migration patterns of several known PSII auxiliary factors differ between the *psb28* mutant and the WT

We next asked whether known PSII auxiliary factors would accumulate in complex with the accumulating early PSII assembly intermediates in the *psb28* mutant. To investigate this, we started out from a list of PSII auxiliary factors compiled by Lu (2016) for Arabidopsis and searched for *Chlamydomonas* homologs that were detected with three replicates each in the mutant and the WT in our CP dataset (Supplemental Table S2). This resulted in 26 factors, all of which overaccumulated in *psb28* compared to the WT, with the exception of SRP43, ALB3.1, TEF30 and LPA2 (Supplemental Table S2). The heat map of the migration profiles in Figure 6B shows that most of the 26 factors migrated in the low molecular mass region below CP43_mod_. Of the factors found in assemblies above CP43_mod_, we found eight to display significant differences in at least one gel band (P < 0.05) between mutant and WT (Figures 6B, 7; Supplemental Figure S11): STL1, FTSH1, FTSH2, TEF30, HCF244, HCF136, TEF5, and PsbN. TEF30 (MET1 in Arabidopsis) interacts with PSII monomers and facilitates PSII supercomplex formation (Bhuiyan et al., 2015; Muranaka et al., 2016). We did not find TEF30 migrating with PSII monomers in the *psb28* mutant, suggesting that there are no PSII monomers capable of binding TEF30 (Figure 6B). STL1 and FTSH1/2 accumulated in the *psb28* mutant above WT levels in several gel bands with higher molecular mass complexes (Supplemental Figure S11). STL1 is homologous to STN8 in Arabidopsis, which phosphorylates PSII core subunits as well as PGRL1-A to regulate cyclic electron flow (CEF) (Bonardi et al., 2005; Reiland et al., 2011). Although STL1 has not been characterized in Chlamydomonas, a role in CEF regulation might be conserved (Longoni and Goldschmidt-Clermont, 2021). The very similar migration pattern in large molecular mass complexes of FTSH1 and FTSH2 (Figure 6B; Supplemental Figure S11) confirms their presence in heterooligomers and their higher abundance in the *psb28* mutant points to a role of this thylakoid membrane protease in degradation of misassembled PSII complexes (Malnoe et al., 2014).

In contrast to STL1 and FTSH1/2, all other PSII auxiliary factors accumulating in complexes above CP43_mod_ at significantly higher levels in the mutant compared with the WT co-migrated with early PSII assembly intermediates: HCF244, HCF136, TEF5, and PsbN with PSII monomers/RC47, and TEF5 and PsbN with RCII (Figure 7). No such peaks were observed for any of the four factors in the *lpa2* mutant (Supplemental Figure S12). There might be some co-migration of PsbN and HCF244 with CP43_mod_ and of HCF136, HCF244, and TEF5 with very small assemblies of D1 and PsbE/F. HCF244 (Ycf39 in *Synechocystis*), HCF136 (Ycf48 in *Synechocystis*), and PsbN have been found in early PSII assembly intermediates with roles in PSII assembly in cyanobacteria and plants (Meurer et al., 1998; Plucken et al., 2002; Komenda et al., 2008; Link et al., 2012; Knoppova et al., 2014; Torabi et al., 2014; Knoppova et al., 2022). In contrast, Arabidopsis PSB33/LIL8 (the TEF5 homolog in land plants) has been found to co-migrate only with RC47 and larger PSII assemblies (especially monomers) (Fristedt et al., 2015; Fristedt et al., 2017; Kato et al., 2017; Nilsson et al., 2020).

**Figure 7.**
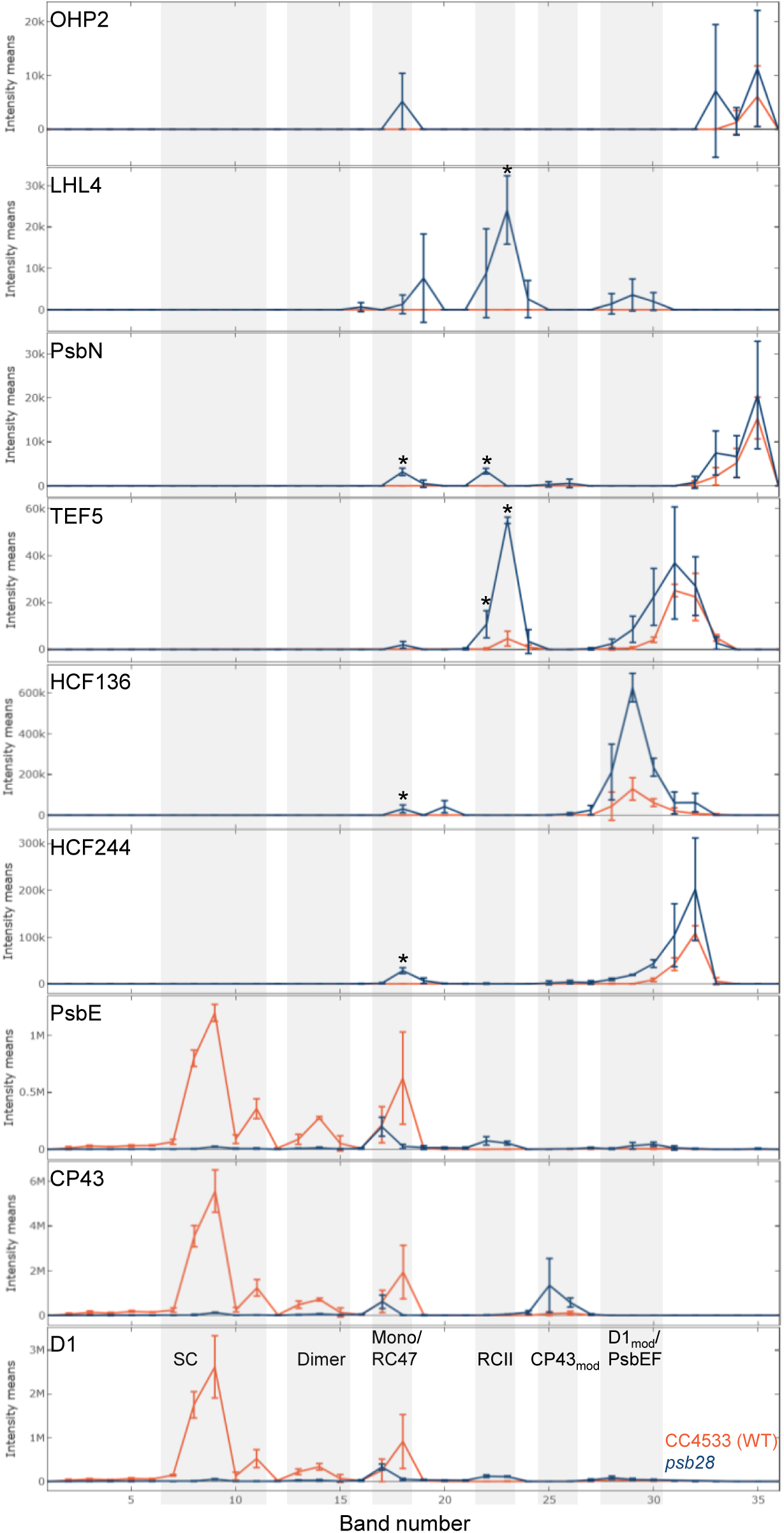
BN-PAGE migration profiles of PSII core subunits and of putative novel PSII-associated proteins. Values for each protein are derived from averaged peptide ion intensities from three biological replicates. Error bars represent standard deviation. Individual profiles from each replicate before and after normalization and statistical analyses can be accessed in Supplemental Dataset S2. Asterisks indicate significant differences in ion intensities between WT (red) and *psb28* mutant (blue) in bands containing complexes larger than CP43_mod_ (two-tailed unpaired t-test, *P* < 0.05). SC – supercomplexes.

### Identification of novel proteins potentially involved in early PSII assembly steps

To identify potential novel factors associated with early PSII assemblies, we searched in our CP dataset for chloroplast proteins with similar migration properties as the four known PSII auxiliary factors HCF244, HCF136, TEF5, and PsbN, i.e., proteins specifically accumulating in bands 17/18 (PSII monomers/RC47) and/or 22/23 (RCII) in *psb28* but not in the WT. Five proteins met these criteria: OHP2 (two peptides), Cre03.g154600 (three peptides), and Cre01.g007700 (three peptides) with peaks in bands 17/18, LHL4 (four peptides) in bands 22/23, and Cre10.g450500 (three peptides) in both (Figure 7; Supplemental Figure S12). For OHP2, Cre01.g007700, and LHL4 no such peaks were observed in the *lpa2* complexome profiling dataset or they were not detected at all. In that dataset, Cre10.g450500 co-migrated with PSII monomers and RC47, while Cre03.g154600 migrated between them, thus disqualifying Cre03.g154600 as a PSII-associated protein. OHP2 and OHP1 together with HCF244 form a complex that has been found to be essential for Chl integration into PSII, or for protection of the newly synthesized Chl-associated D1 during formation of RCII in Arabidopsis (Hey and Grimm, 2018; Li et al., 2018; Myouga et al., 2018). In the Chlamydomonas *ohp2* knockout mutant newly made D1 is rapidly degraded resulting in a complete lack of PSII (Wang et al., 2023). LHL4 is an LHC-like protein closely related to PSBS and uniquely found in green microalgae (Dannay et al., 2024). Cre01.g007700 encodes an aminopeptidase and Cre10.g450500 has a starch-binding domain, both are yet uncharacterized in *Chlamydomonas*.

### The *tef5* mutant has a lower PSII content than the WT and shows impaired growth in high light and under photoautotrophic conditions

The co-migration of a large portion of TEF5 with RCII in the *psb28* mutant suggested a possible role of TEF5 in PSII biogenesis, which was considered unlikely for its homolog PSB33/LIL8 in Arabidopsis (Fristedt et al., 2017; Kato et al., 2017). TEF5/PSB33/LIL8 are conserved in the green lineage and there are no orthologs in cyanobacteria. They contain chloroplast transit peptides and share a Rieske-like domain lacking the residues required for the binding of mononuclear iron or an iron-sulfur cluster (Fristedt et al., 2015) (Figure 8A). Moreover, they share one to two C-terminal transmembrane helices, where the loss of one transmembrane helix appears to have occurred before the evolution of land plants (Chlamydomonas and *Ostreococcus tauri* contain two, while *Chlorella variabilis* and members of the Streptophytes contain only a single transmembrane helix, Figure 8A). Although the Rieske-like domains of *Chlamydomonas* TEF5 and Arabidopsis PSB33/LIL8 share only 54% identical residues, their predicted structures are very similar (RMSD = 0.99 Å, TM score = 0.9; Figure 8B). To investigate a possible role of TEF5 in early steps of PSII assembly, we selected a *tef5* mutant from the CLiP collection (Li et al., 2016) that had the CIB1 mutagenesis cassette integrated into the sixth intron of the *TEF5* gene (Figure 8C). Since we could amplify *TEF5* sequences of the expected sizes from both sides of the CIB1 cassette by PCR, there appear to be no larger deletions/rearrangements (Supplemental Figure S13A, B). qRT-PCR analysis revealed a ∼147-fold reduced abundance of *TEF5* transcript in the *tef5* mutant compared with the WT (Supplemental Figure S13C). An antibody raised against a peptide from the N-terminal part of the TEF5 protein (Figure 8A) specifically detected a protein band at the expected molecular mass of ∼27.5 kDa in the WT, which was absent in the *tef5* CLiP mutant (Figure 8D; Supplemental Figure S13D). In the *tef5* mutant, PSII core subunits accumulated to between 20% and 40% of WT levels while there was no or little change in the abundance of LHCII, PSI core subunits, ATP synthase subunit CF1β, and Cyt *f* (Figures 8D, E). We synthesized the *TEF5* cDNA sequence interrupted by the first two *Chlamydomonas RBCS2* introns, fused it with sequences encoding a C-terminal 3xHA tag or multiple stop codons and placed it under control of the constitutive *HSP70A-RBCS2* promoter and the *RPL23* terminator using Modular Cloning (Figure 8C). We combined the *TEF5* transcription unit with an *aadA* cassette and transformed it into the *tef5* mutant. Spectinomycin resistant transformants obtained with both constructs were then screened by immunoblotting for the accumulation of HA-tagged TEF5 and/or for enhanced D1 accumulation (Supplemental Figure S14). Five transformants accumulated HA-tagged TEF5 and all accumulated D1 to WT levels. One transformant (*tef5*-cHA) accumulated TEF5 transcripts to ∼73-fold higher levels than WT but TEF5 protein levels were not much higher than those in the WT (Figure 8D; Supplemental Figure 13D). Since the band detected with the TEF5 antibody in cHA had the same size as in WT, we assume that the 3xHA tag was removed from part of the protein, as was observed with PSB28-3xHA. A transformant generated with the construct encoding non-tagged TEF5 (*tef5*-c15) accumulated TEF5 to much higher levels than the WT (Figure 8D). Both, HA-tagged and untagged transformants accumulated PSII subunits to WT levels and perhaps even beyond (Figure 8D, E; Supplemental Figure S14B, C). This correlated with the restoration of WT Fv/Fm values (Figure 8F). The *tef5* mutant showed slightly reduced growth under photoautotrophic and mixotrophic conditions in low light (30 µmol photons m^-2^ s^-1^), and a severe growth defect under mixotrophic conditions in high light (600 µmol photons m^-2^ s^-1^), while growth under heterotrophic conditions in the dark was like WT (Figure 8G). These growth phenotypes were fully restored in the *tef5*-c15 transformant.

**Figure 8.**
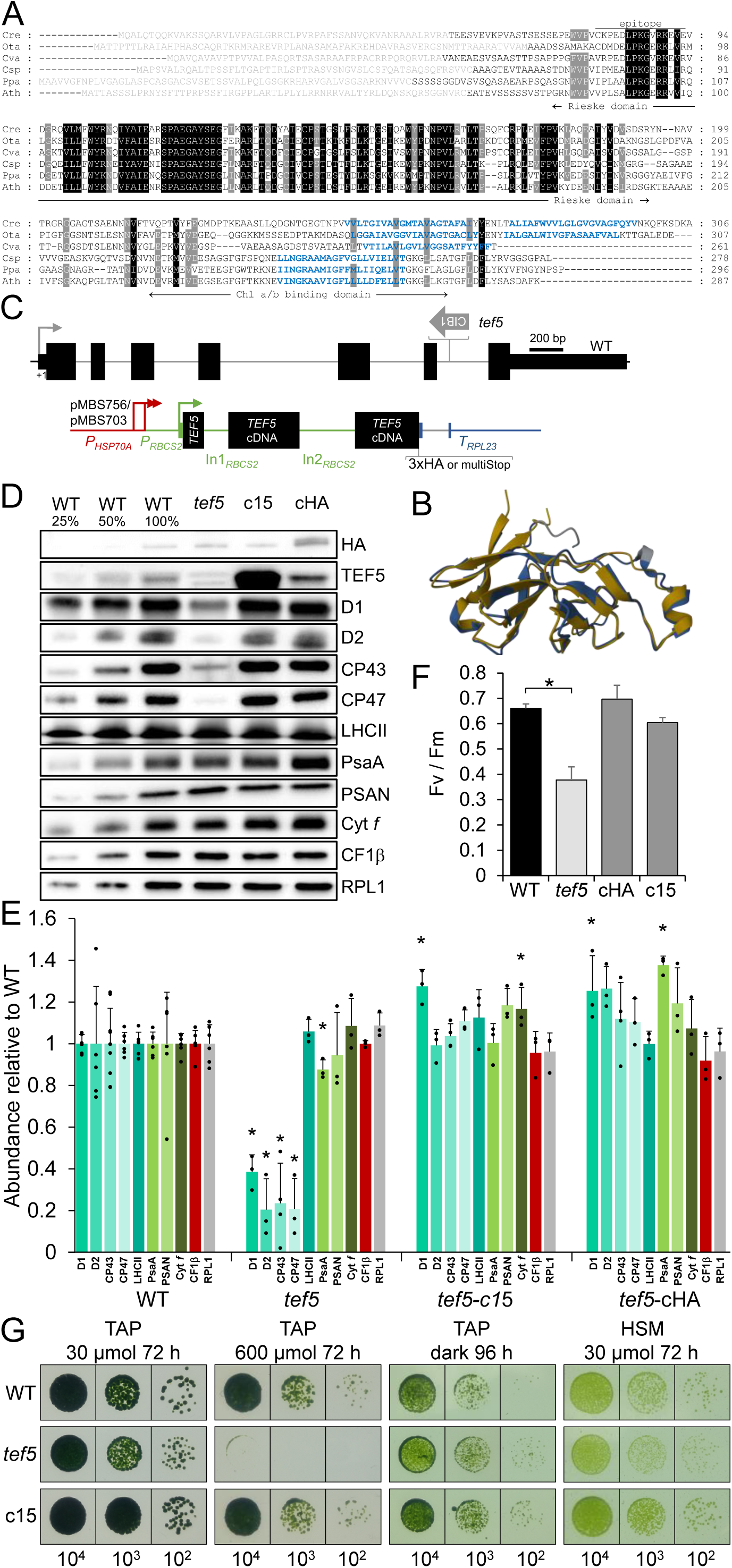
Phenotypes of the *tef5* mutant compared to WT and complemented lines. **(A)** Alignment of amino acid sequences of algal and land plant homologs of TEF5/PSB33/LIL8. Residues highlighted in black and gray are conserved in six and five of the sequences, respectively. Predicted chloroplast transit peptides are shown in gray, predicted transmembrane helices in blue. The epitope from *Chlamydomonas* TEF5 used for antibody production is indicated by a horizontal line. Cre – *Chlamydomonas reinhardtii* (Cre09.g411200), Ota – *Ostreococcus tauri* (XP_003078526), Cva – *Chlorellla variabilis* (XP_005846469), Csp – *Closterium sp*. (CAI5958768), Ppa – *Physcomitrium patens* (XP_024377109), Ath – *Arabidopsis thaliana* (AT1G71500). **(B)** Pairwise structure alignment of the Rieske-like domains from Arabidopsis PSB33 (gold) and *Chlamydomonas* TEF5 (blue). **(C)** Structure of the *Chlamydomonas TEF5* gene, insertion site of the CIB1 cassette in the *tef5* mutant, and constructs for complementation. Protein coding regions are drawn as black boxes, untranslated regions as bars, and introns (In) and promoter regions as thin lines. Arrows indicate transcriptional start sites. **(D)** Immunoblot analysis of the accumulation of TEF5 and of subunits of the major thylakoid membrane protein complexes. c15 and cHA are lines complemented with constructs pMBS703 and pMBS756, respectively, shown in (C). PSII – D1, D2, CP43, CP47, LHCII; PSI – PsaA, PSAN; Cyt *b_6_f* complex – Cyt *f*; ATP synthase – CF1b. Ribosomal protein RPL1 served as loading control. 10 µg of whole-cell proteins (100%) were analysed. **(E)** Quantification of the immunoblot analysis shown in (D). Values are means from three independent experiments normalized first by the median of all signals obtained with a particular antiserum in the same experiment, and then by the mean signal of the WT. Error bars represent standard deviation. Asterisks indicate significant differences with respect to the WT (two-tailed, unpaired *t*-test with Bonferroni-Holm correction, *P* < 0.05). The absence of an asterisk means that there were no significant differences. **(F)** F_v_/F_m_ values of the *tef5* mutant versus WT and complemented lines. Shown are averages from three to seven independent experiments each measured with three technical replicates. Error bars represent standard deviation. Asterisks indicate significant differences with respect to the WT (two-tailed, unpaired *t*-test with Bonferroni-Holm correction, *P* < 0.001). The absence of an asterisk means that there were no significant differences. **(G)** Analysis of the growth of 10^4^ – 10^2^ spotted cells under the conditions indicated.

### Chloroplast morphology is intact in the *tef5* mutant, but thylakoid membranes are swollen

Light and transmission electron microscopy (TEM) were used to analyze possible changes in cell morphology and thylakoid ultrastructure in the *tef5* mutant. Light microscopy revealed no visible change in chloroplast morphology in the *tef5* mutant (Figure 9A). TEM revealed that thylakoid membranes in the mutant are more loosely packed and swollen (Figure 9B).

**Figure 9.**
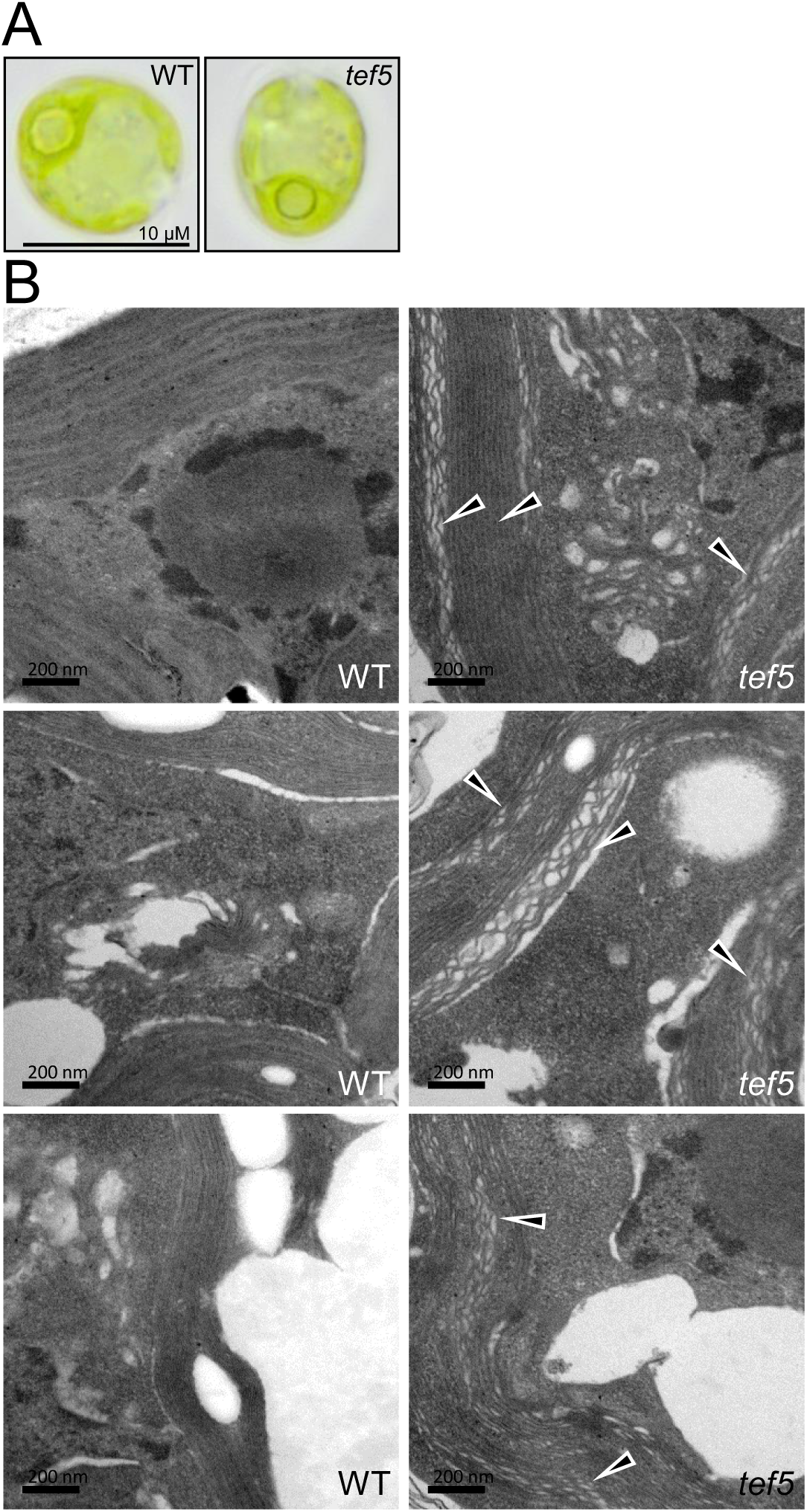
Light and electron microscopy of the *tef5* mutant. **(A)** Light microscopy images of WT and *tef5* mutant grown under mixotrophic conditions in low light (30 µmol photons m^-2^ s^-1^). **(B)** Electron microscopy pictures of WT (left) and *tef5* mutant (right) grown under mixotrophic conditions in low light. Black triangles indicate swollen thylakoids in the mutant.

### The synthesis of CP47 and PsbH is reduced in the *tef5* mutant

To investigate whether the absence of TEF5 affects the synthesis of PSII core subunits in the *tef5* mutant, we performed pulse-chase analyses with ^14^C-acetate. As judged from the ^14^C-labeling of proteins within the 7-min ^14^C-acetate pulse, synthesis of CP47 and PsbH appeared reduced in the *tef5* mutant when compared with the WT. Based on the 20-min chase period, D1 appeared less stable in the *tef5* mutant (Figure 10). Both phenotypes were restored to WT in the complemented lines.

**Figure 10.**
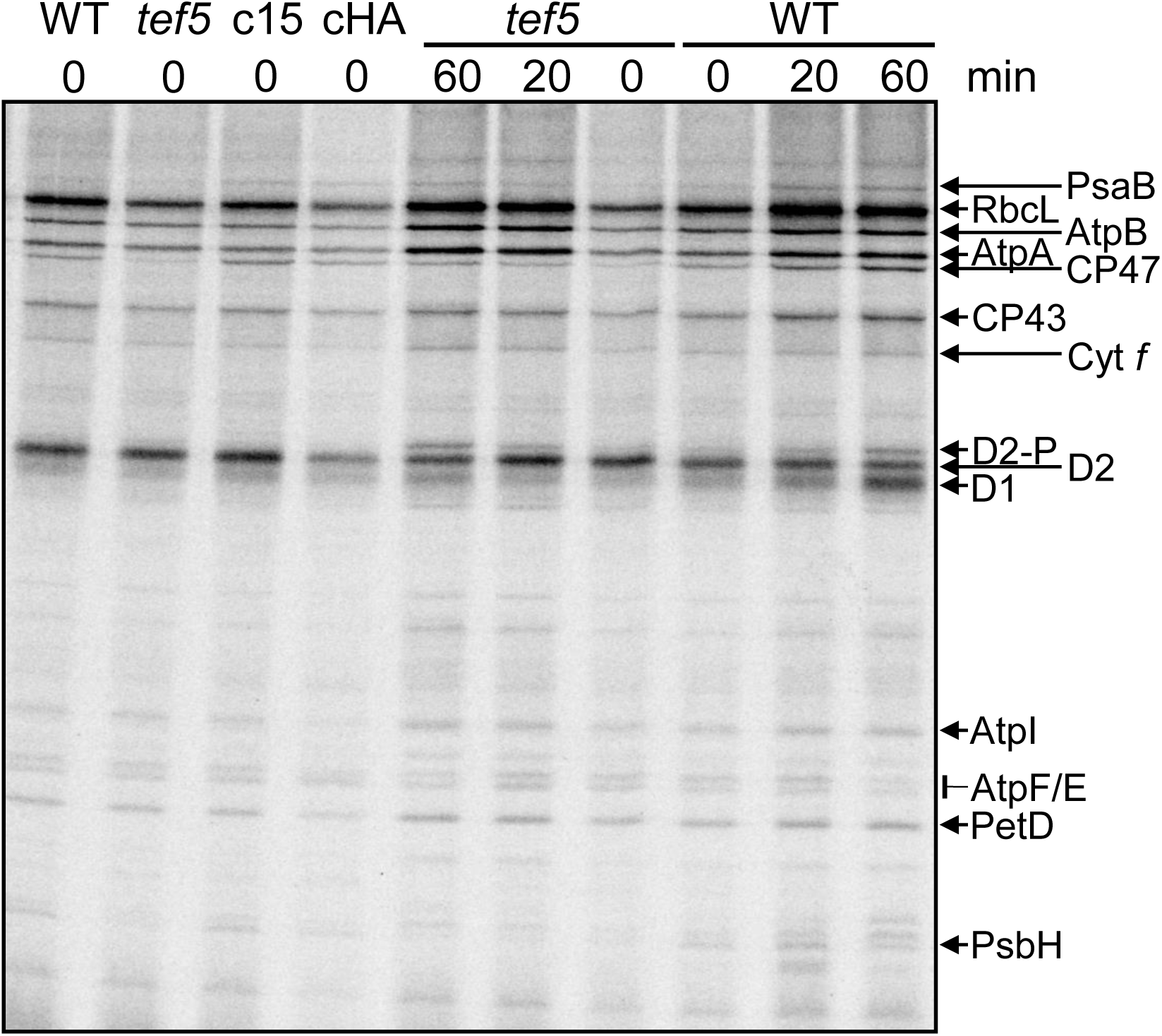
Pulse-chase analysis of synthesis and stability of thylakoid membrane proteins in the *tef5* mutant. WT, *tef5* mutant and complemented lines c15 and cHA were labelled with ^14^C-acetate in low light (20 µmol photons m^-2^ s^-1^) for 7 min in the presence of cytosolic translation inhibitor cycloheximide (0) and chased with unlabelled acetate for 20 and 60 min. Proteins were separated on a 12-18% SDS-urea gel and visualized by autoradiography.

### PSII assembly is impaired in the *tef5* mutant in the light but to a lesser extent in the dark

To assess how the reduced synthesis and accumulation of PSII core subunits in the *tef5* mutant affects PSII complex assembly, we analyzed whole-cell proteins from the low light-grown WT, the *tef5* mutant, and the complemented line *tef5*-c15 by BN-PAGE and immunoblotting using antibodies against D1 and CP43. As shown in Figure 11A, we found much weaker signals for PSII supercomplexes, dimers, and monomers in the mutant when compared with WT and the complemented line. In contrast, the *tef5* mutant accumulated RCII and CP43_mod_, which was not the case in the WT and the complemented line.

**Figure 11.**
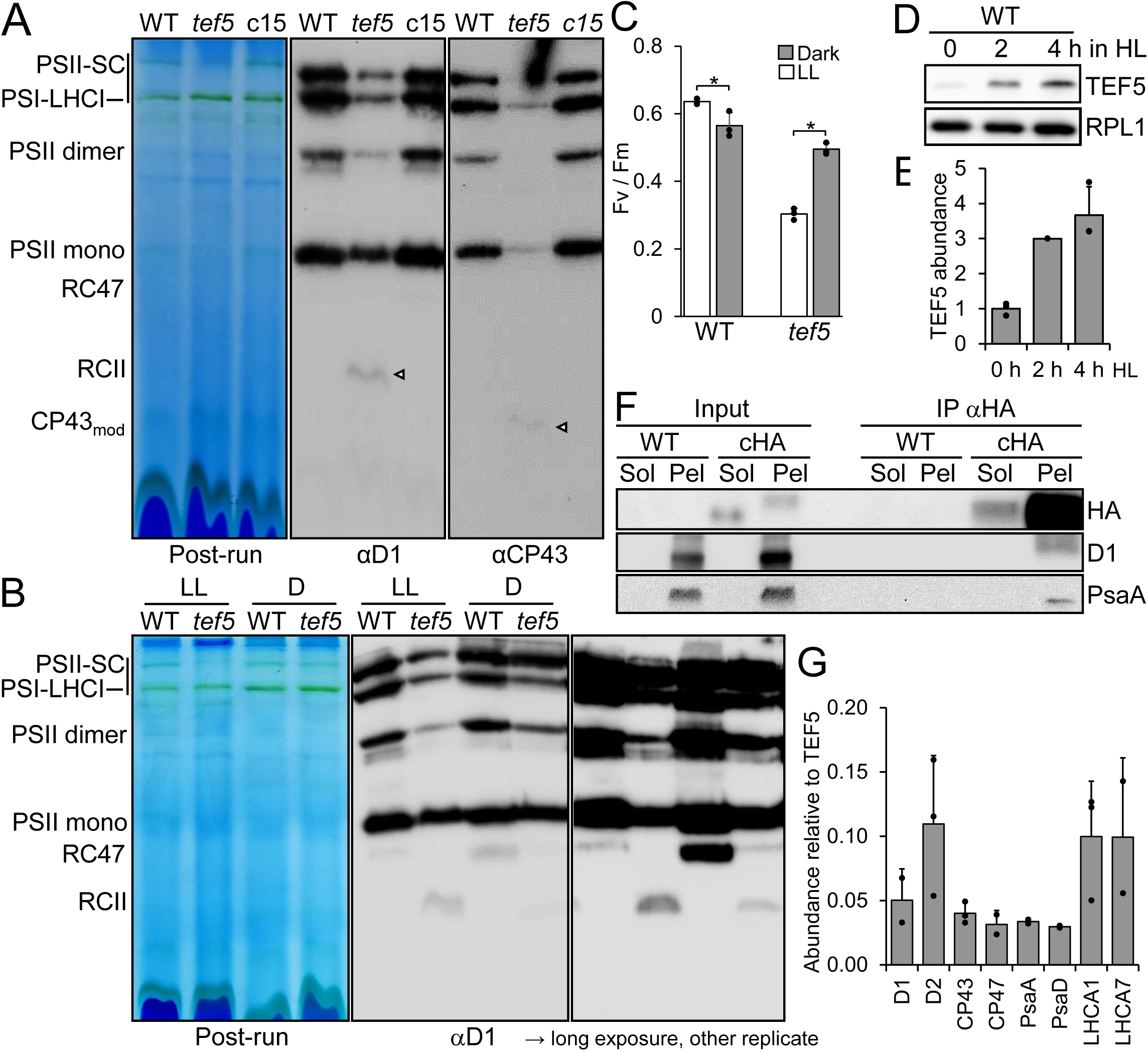
Analysis of protein complexes in the *tef5* mutant and of proteins interacting with TEF5. **(A)** BN-PAGE analysis of proteins from cells grown in low light (30 µmol photons m^-2^ s^-1^). 60 µg of whole-cell proteins from WT, *tef5* mutant, and complemented line *tef5*-c15 were solubilized with 1% β-DDM and separated on a 4-15% BN gel. Shown is a picture of the gel after the run and an immunoblot detected with antibodies against D1 and CP43. Arrowheads point to faint bands likely representing RCII and CP43_mod_ in the *tef5* mutant. SC – supercomplexes. **(B)** BN-PAGE analysis of proteins from WT and *tef5* mutant grown in low light (LL, 30 µmol photons m^-2^ s^-1^) and in the dark (D) for 72 h. Whole-cell proteins were solubilized with 1% β-DDM and separated on a 4-15% BN gel. Shown is a picture of the gel after the run and an immunoblot detected with an antibody against D1 accompanied by a longer exposure of an independent replicate. **(C)** F_v_/F_m_ values of the *tef5* mutant versus WT grown in low light light (LL, 30 µmol photons m^-2^ s^-1^) and in the dark for 72 h. Shown are averages from three independent experiments. Error bars represent standard deviation. Asterisks indicate significant differences between low-light versus dark-grown cells (two-tailed, unpaired *t*-test, *P* < 0.05.). **(D)** Analysis of TEF5 accumulation in high light (HL). WT was exposed to 1200 µmol photons m^-2^ s^-1^ for 4 h and samples taken prior, 2 and 4 h after the treatment were analysed by immunoblotting using the peptide antibody against TEF5 and an antibody against RPL1 as loading control. **(E)** Quantification of the immunoblot analysis shown in (D). Values are means from three independent experiments. Normalization was done as described for Figure 1D. **(F)** Immunoprecipitation of TEF5. Cells from complemented line *tef5*-cHA were fractionated via freeze-thaw cycles and centrifugation. HA-tagged TEF5 was then immunoprecipitated (IP) from soluble (Sol) and membrane-enriched (Pel) fractions with an HA antibody. 1% of the input and 10% of the precipitate were analysed by SDS-PAGE and immunoblotting using antibodies against HA, D1, and PsaA. **(G)** Mass spectrometry-based quantification of PSI and PSII subunits co-precipitated from solubilized membrane fractions with HA-tagged TEF5. IBAQ values for each protein were normalized by the IBAQ value for TEF5. Shown are mean values from 2-3 independent replicates. Error bars represent standard deviation.

To investigate whether the impaired assembly of PSII in the *tef5* mutant was due to an effect of light, we compared protein complexes in solubilized whole-cell extracts from the WT and the *tef5* mutant grown in low light and in the dark for 72 h by BN-PAGE and immunoblotting using a D1 antibody. We clearly observed stronger signals for PSII dimers and supercomplexes in the dark-versus light-grown *tef5* mutant (Figure 11B). Moreover, the mutant accumulated less RCII in the dark than in the light. Most interestingly, the mutant accumulated no RC47 in the light. In the dark, the WT accumulated large amounts of RC47 and the *tef5* mutant accumulated some. The partially rescued PSII assembly in the dark-grown mutant was also reflected at the level of Fv/Fmax values, which were significantly higher in the dark-versus low light-grown mutant (0.5 vs.0.3, P < 0.001) but did not fully reach values obtained for the dark-grown WT (0.57) (Figure 11C).

The potential role of TEF5 in PSII assembly at the RCII/RC47 level suggests that TEF5 may also be involved in the repair of photodamaged PSII. We therefore first tested whether TEF5 accumulates in cells exposed to high light and found a ∼3.7-fold increased abundance of TEF5 protein after 4 h exposure to 1200 µmol photons m^-2^ s^-1^ (Figure 11D, E). This was surprising, since PSB33/LILI8 was reported to be expressed constitutively (Kato et al., 2017). To investigate the susceptibility of PSII in the *tef5* mutant to high light and its capability to recover functional PSII, we exposed the WT, the *tef5* mutant, and the complemented lines to high light (1800 µmol photons m^-2^ s^-1^) in the presence of CAP for one hour and allowed cells to recover in the presence and absence of CAP at low light (30 µmol photons m^-2^ s^-1^).

All four lines recovered 86-96% of initial PSII activity (and most of D1) at similar rates within 6.5 hours in a protein synthesis-dependent manner (Supplemental Figure S15). Like the *psb28* mutant, the *tef5* mutant lost PSII activity upon sulfur starvation faster than the WT and the complemented lines but recovered initial PSII activity (and D1 levels) with similar rates as the other lines (Supplemental Figure S16). In summary, the *tef5* mutant is impaired in PSII assembly presumably at the step where the CP47_mod_ combines with RCII to RC47. As observed for the *psb28* mutant, the low levels of PSII made in the *tef5* mutant are susceptible to photoinhibition and degradation upon sulfur deprivation but can be fully recovered to these low levels at WT rates.

### TEF5 interacts with subunits of PSI and PSII

The co-migration of TEF5 with early PSII assembly intermediates in the *psb28* mutant and its potential role in PSII assembly implies its direct interaction with PSII. To test this, we used the HA antibody to immunoprecipitate TEF5-3xHA from soluble and membrane-enriched fractions from the complemented *tef5*-cHA line. Prior to immunoprecipitation, complexes were stabilized by *in-vivo* crosslinking with 0.37% formaldehyde. As shown in Figure 11F, most of TEF5-3xHA was precipitated from the membrane-enriched fraction but some was also precipitated from the soluble fraction. The different migration behavior of TEF5 in soluble and membrane-enriched fractions could be due to the presence of large amounts of LHCII at the same position in the gel only in the membrane-enriched fraction, which has been observed also for Arabidopsis PSB33/LIL8 (Kato et al., 2017). D1 and PsaA were co-precipitated with TEF5 only in the membrane-enriched fraction. To identify and quantify all proteins interacting with TEF5, we analyzed the TEF5 immunopreciptates by LC-MS/MS. In line with the immunoblot data we detected ∼16 times more TEF5 in the membrane-enriched fraction than in the soluble fraction (Supplemental Dataset S4). Among the proteins detected with TEF5 in at least two replicates in the membrane fraction, we found PSII subunits D1, D2, CP43, and CP47 as well as PSI subunits PsaA, PsaD, LHCA1, and LHCA7 (Figure 11G; Supplemental Dataset S4). We also detected RBCS2, a transporter and an ATPase subunit from mitochondria, and a putative transhydrogenase, guanylate cyclase, and nucleolar protein which most likely are contaminants. Intensity-based absolute quantification (IBAQ) normalized to TEF5 revealed that D2 is the most prominent TEF5 interaction partner, followed by LHCA1/7, D1, CP43, PsaA, CP47, and PsaD (Figure 11G).

## Discussion

### Psb28 is of much greater importance for PSII assembly in Chlamydomonas than in Synechocystis

Chlamydomonas PSB28 has several traits in common with cyanobacterial Psb28. Cyanobacterial Psb28 is sub-stoichiometric to PSII and interacts only transiently with PSII, mainly with RC47 and less with PSII monomers, while most Psb28 is present as free protein (Kashino et al., 2002; Dobakova et al., 2009; Boehm et al., 2012; Nowaczyk et al., 2012; Sakata et al., 2013; Beckova et al., 2017; Xiao et al., 2021; Zabret et al., 2021). Similarly, Chlamydomonas PSB28 is ∼78-fold less abundant than PSII and in complexome profiling was only found as free protein in WT, but co-migrated mainly with RC47 and less with PSII monomers in the *lpa2* mutant which overaccumulates RC47 (Spaniol et al., 2022). HA-tagged PSB28 co-migrated more with PSII monomers than with RC47 and a fraction of tagged PSB28 was present as free protein (Figure 4A). Accordingly, immunoprecipitation of tagged PSB28 revealed D2 and D1 as the most prominent interaction partners, followed by CP47 and CP43 (Figure 4B, C). Functional similarity between Chlamydomonas and cyanobacterial Psb28 was indicated by their structural similarity (Figure 1B) and by the ability of *Synechocystis* Psb28-1 to partially complement the Chlamydomonas *psb28* mutant (Figure 5). Common is the reduced synthesis of CP47 and PSI/PsaB in *Synechocystis* and Chlamydomonas *psb28* mutants (Dobakova et al., 2009; Beckova et al., 2017), however, in Chlamydomonas the synthesis of D1, D2, CP43, and PsbH was affected, too (Figure 3). This points to control by epistasis of synthesis (CES) of PSII subunits in the *psb28* mutant (Minai et al., 2006), possibly by a negative feedback regulation effected by accumulating assembly intermediates such as RCII and CP43_mod_ (Figures 4A, 6A; Table 2).

One difference between *Synechocystis* Psb28-1 and Chlamydomonas PSB28 is that the abundance of Psb28-1 did not increase at high light intensities (Beckova et al., 2017), while the abundance of PSB28 increased ∼2.9-fold (Figure 4E, F). Moreover, *Synechocystis* Psb28-1 and 2 were found in PSII-PSI supercomplexes particularly under high light intensities (Beckova et al., 2017), which we did not observe in Chlamydomonas (Figure 4D). In Chlamydomonas, more PSB28 interacted with PSII monomers and particularly with RC47 in high versus low light (Figure 4D), suggesting a role of PSB28 also in PSII repair in this alga.

Probably most surprising are the differences in the phenotypes of the *Synechocystis* and Chlamydomonas *psb28* mutants: the *Synechocystis psb28-1* mutant accumulated fully functional PSII and growth phenotypes were observed only at higher temperatures and high or fluctuating light exposure (Dobakova et al., 2009; Sakata et al., 2013; Beckova et al., 2017). In contrast, the Chlamydomonas *psb28* mutant could not grow photoautotrophically (Figure 1H) and accumulated PSII supercomplexes, dimers and monomers only to 1%, 6% and 27% of WT levels, respectively, while it overaccumulated RCII and CP43_mod_ (Table 2; Figures 4A, 5D, 6A). PSII outer antennae were reduced by ∼45% and PSI/LHCI by 16-19%, compared with WT. Levels of the ATP synthase were unaltered in the mutant, while the Cyt *b_6_f* complex overaccumulated between 1.4- and 1.9-fold (Table 2, Figure 1D, E). These dramatic changes in the photosynthetic apparatus probably cause reduced thylakoid stacking and a distorted shape of the chloroplast (Figure 2). The reduced levels of PSI, increased levels of Cyt *b_6_f* and the distorted chloroplast shape are unusual phenotypes for PSII mutants in Chlamydomonas: The *ohp2* mutant, lacking PSII, accumulates PSI and Cyt *b_6_f* at WT levels (Wang et al., 2023) as do the *lpa2* and *tef5* mutants, with PSII levels reduced by about half and below 40% of the WT levels, respectively (Spaniol et al., 2022) (Figure 8D, E). While some changes in thylakoid structure were observed in the *lpa2* and *tef5* mutants, the morphology of the chloroplast was unaltered (Spaniol et al., 2022) (Figure 9). The reason for these pleiotropic phenotypes of the *psb28* mutant could be an additional function of PSB28 besides that as a PSII assembly factor. Indeed, *Synechocystis* Psb28-1 was proposed to play a role in regulating chlorophyll incorporation into CP47 and PSI (Beckova et al., 2017). Alternatively, PSII assembly intermediates specifically accumulating in the *psb28* mutant might act as regulators of other processes resembling CES (Choquet and Wollman, 2023). It is also possible that PSII intermediates in the *psb28* mutant specifically bind assembly factors, chaperones or proteases, which are then not sufficiently available for other chloroplast processes.

The fact that PSII accumulation is much more affected in the Chlamydomonas *psb28* mutant than in the *Synechocystis psb28* mutant is probably due to the very efficient proteolytic degradation of non-assembled complex subunits in Chlamydomonas (Choquet and Wollman, 2023). Accordingly, we observed a rapid removal particularly of newly synthesized D2 in the *psb28* mutant (Figure 3). Moreover, it is likely that also misassembled complexes are subject of efficient proteolytic degradation in Chlamydomonas, possibly explaining why the Chlamydomonas *psb28* mutant barely accumulated any larger PSII assemblies in contrast to the *Synechocystis psb28* mutant. Degradation of unstable PSII assemblies was also proposed for the Chlamydomonas *lpa2* mutant (Spaniol et al., 2022). A high proteolytic activity in Chlamydomonas is also indicated by the partial removal of the 3xHA tag from the PSB28- and TEF5-3xHA fusion proteins (Figures 1D, 8D; Supplemental Figures S2, S14B). In line with this idea, FTSH1/2 were ∼2.5-fold more abundant in *psb28* than in WT (Supplemental Table S2) and FTSH1/2 complexes were more abundant in the higher molecular mass range (Figure 6B; Supplemental Figure S11). Another explanation for the impaired accumulation of larger PSII assemblies in the Chlamydomonas *psb28* mutant is that the conformational changes introduced by Psb28 into the PSII core (Xiao et al., 2021; Zabret et al., 2021) are more important for correct assembly of the CP43_mod_ into RCII in Chlamydomonas than in *Synechocystis*. It was proposed that the conformational changes introduced by Psb28 might also protect premature PSII from photodamage (Xiao et al., 2021; Zabret et al., 2021). Since the problem in PSII assembly prevailed in the dark-grown Chlamydomonas *psb28* mutant (Supplementary Figure S4A), the impaired accumulation of larger PSII assemblies in the mutant is unlikely to be caused by enhanced photodamage to early PSII assemblies.

The dependence of PSII assembly on auxiliary factors generally appears to be stronger in chloroplasts than in cyanobacteria. Examples for this, in addition to PSB28, are HCF136 (YCF48 in cyanobacteria), HCF244 (Ycf39 in cyanobacteria), PsbN, and PAM68. While the absence of these factors resulted in severe PSII assembly defects in Arabidopsis or tobacco, *Synechocystis* mutants lacking these factors could assemble functional PSII (Mayers et al., 1993; Meurer et al., 1998; Komenda et al., 2008; Armbruster et al., 2010; Link et al., 2012; Knoppova et al., 2014; Torabi et al., 2014).

### Complexome profiling confirms PBA1 and CGLD16 as potential novel PSII-associated proteins

Previously, complexome profiling of thylakoid membranes of the WT and the *lpa2* mutant identified PBA1 (putatively Photosystem B Associated 1) and CGLD16 as potential novel PSII-associated proteins (Spaniol et al., 2022). Both contain predicted single transmembrane helices and chloroplast transit peptides and have predicted mature masses of 6.4 and 7.9 kDa, respectively. PBA1 is present only in members of the green algae, brown algae, diatoms, and Eustigmatophytes, while CGLD16 is conserved in the green lineage and diatoms. CGLD16 co-migrated with PSII monomers and RC47 in the WT and the *lpa2* mutant (Spaniol et al., 2022), and we found the same migration pattern for CGLD16 also for the WT and the *psb28* mutant in this work (Supplemental Figure S12). In the WT, PBA1 co-migrated with PSII supercomplexes, dimers, monomers, and RC47 and its abundance in these complexes was reduced in the *lpa2* mutant, where the unassembled form was more abundant (Spaniol et al., 2022). In this work, we found PBA1 to co-migrate with PSII supercomplexes only in WT and with PSII monomers/RC47 in WT and *psb28* (Figure 6A; Supplemental Figure S10). These data confirm that PBA1 and CGLD16 might be novel PSII-associated proteins. Cryo-EM analyses of PSII from Chlamydomonas have revealed two new densities referred to as unidentified stromal protein (USP) and small luminal protein (SLP) (Sheng et al., 2019; Sheng et al., 2021). Perhaps these densities are attributable to CGLD16 and PBA1? Nonetheless, these structural studies show that not all of the PSII-associated proteins have been already discovered, at least not in Chlamydomonas.

### Complexome profiling confirms previously identified PSII assembly factors and identifies new factors with potential roles in early PSII assembly

Complexome profiling of the thylakoid membranes of WT and *psb28* revealed 26 PSII auxiliary factors known from previous studies (Lu, 2016) (Supplemental Table 2). Among these, 22 accumulated to higher levels in the mutant compared to WT, potentially to compensate for impaired PSII accumulation in the mutant. Six proteins were found to co-migrate with early PSII assembly intermediates (monomers/RC47 and smaller) specifically in *psb28* but not in *lpa2* or WT (Figure 6B, 7; Supplemental Figure S12). These were PsbN, HCF136, HCF244, OHP2, TEF5, and LHL4. Roles in early PSII assembly steps have been reported for PsbN, HCF136, HCF244, and OHP2, which forms a complex with HCF244 and OHP1 (Meurer et al., 1998; Plucken et al., 2002; Komenda et al., 2008; Link et al., 2012; Knoppova et al., 2014; Torabi et al., 2014; Knoppova et al., 2022; Wang et al., 2023). As discussed below, our data indicate a role also for TEF5 in early PSII assembly. Overall, this highlights the power of the complexome profiling approach to identify assembly factors that enrich with assembly intermediates in assembly mutants (Heide et al., 2012; Spaniol et al., 2022).

Only LHL4 was not assigned a role in PSII assembly. LHL4 is an LHC-like protein harboring three transmembrane domains of which the region around the first transmembrane helix shares high sequence similarity with the same region in PSBS and with cyanobacterial HliA-D (Supplemental Figure S17) (Dannay et al., 2024). LHL4 is uniquely found in green microalgae, and in Chlamydomonas the *LHL4* gene is induced upon UV-B and high light treatment (Teramoto et al., 2004; Teramoto et al., 2006; Dannay et al., 2024). LHL4 was found to interact with PSII monomers via CP43 and CP47 with a role in protecting PSII from photodamage. LHL4 is barely expressed under low light conditions and interacted with PSII only in the presence of UV-B light (Dannay et al., 2024). We were only able to detect LHL4 in thylakoid membranes of the *psb28* mutant, but not in membranes of the WT or the *lpa2* mutant (Figure 6B, 7; Supplemental Figure S12) (Spaniol et al., 2022). Hence, LHL4 present in low light conditions appears to specifically attach to accumulating early PSII assembly intermediates in the *psb28* mutant, such as RCII and D1_mod_ that did not accumulate in the *lpa2* mutant. Alternatively, PSII assembly intermediates accumulating in the *psb28* mutant might trigger the upregulation of LHL4 (e.g. via enhanced ROS production) that then attaches to the present PSII assemblies. Most likely, LHL4 protects these early PSII assembly intermediates from photodamage or plays a role in binding Chl released from degrading early PSII assemblies, as was proposed for HliC/D in cyanobacteria (Knoppova et al., 2014; Staleva et al., 2015; Knoppova et al., 2022). Since PSII appears to accumulate normally in the *lhl4* mutant (Dannay et al., 2024), it is unlikely that LHL4 plays an essential role during PSII assembly.

### TEF5 is involved in PSII assembly in Chlamydomonas, possibly by facilitating the correct incorporation of the CP47_mod_ into RCII

Features shared by TEF5 and PSB33/LIL8 are the structural similarity of their Rieske-like domains (Figure 8B), and their ability to interact with PSII and PSI (Fristedt et al., 2015; Fristedt et al., 2017; Kato et al., 2017) (Figure 11F, G). Moreover, Chlamydomonas *tef5* and Arabidopsis *psb33/lil8* mutants share a PSII phenotype with reduced accumulation of PSII core subunits, which is constitutive in Chlamydomonas but emerges only under certain environmental conditions in Arabidopsis (Fristedt et al., 2015; Cruz et al., 2016; Fristedt et al., 2017; Nilsson et al., 2020) (Figure 8D, E, G). Arabidopsis *psb33/lil8* showed swollen thylakoids in blue light resembling those of Chlamydomonas *tef5* grown in low light (Figure 9B) (Nilsson et al., 2020). In the Chlamydomonas *tef5* mutant, in low light, PSII subunits accumulated to 20-40% of WT levels (Fig. 8D, E), with monomers, dimers, and supercomplexes accumulating at a lower level than WT and RC47 being undetectable, whereas RCII and CP43_mod_ overaccumulated (Figure 11A, B). The PSII phenotype was attenuated when the *tef5* mutant was grown in the dark, with PSII monomers, dimers and supercomplexes accumulating at higher levels and RCII at a lower level than in the light-grown mutant, and RC47 became detectable (Figure 11B).

As in the Chlamydomonas *tef5* mutant, RC47 was almost undetectable in the *Synechocystis psb28* mutant (Dobakova et al., 2009; Beckova et al., 2017) (Figure 11B), pointing to a role of TEF5 and Psb28 in stabilizing the transient accumulation of RC47. We hypothesize that TEF5 might prime RCII in a way that a correct incorporation of the CP47_mod_ can occur, similar to a possible priming of RC47 by Psb28 to facilitate correct incorporation of the CP43_mod_ (Xiao et al., 2021; Zabret et al., 2021). Consistent with this idea, TEF5 co-migrated with RCII in the WT and to a much greater extent in the *psb28* mutant (Figures 6B, 7). The strong accumulation of RC47 in dark-versus light-grown WT cells points to a slower PSII monomer assembly pace in the dark, which might facilitate correct CP47_mod_ incorporation into RCII even in the absence of TEF5 and would thus explain the attenuated PSII phenotype in the dark-grown *tef5* mutant (Figure 11B). We propose that in the absence of TEF5/PSB33 a fraction of PSII is misassembled, which may result in defects such as a damaged Q_B_ site, as observed by Cruz et al. (2016). An even more effective protein quality control system in Chlamydomonas than in Arabidopsis could explain why such misassembled PSII cores are cleared in Chlamydomonas, whereas they can persist in Arabidopsis. This could be similar in *psb28* mutants. The mild, pale-green phenotype of a rice *psb28* knockout line suggests that some functional PSII can be assembled in the absence of PSB28 (Jung et al., 2008), whereas in the Chlamydomonas *psb28* mutant hardly any functional PSII is made. Here it is possible that the accumulating RCII and CP43_mod_ in the Chlamydomonas *tef5* and *psb28* mutants (Figures 4A; 11A, B) are a mixture of degradation products and assembly intermediates.

To get an idea on how TEF5/PSB33 could interact with PSII, we modelled the structures of Chlamydomonas and Arabidopsis PSII cores comprising D1, D2, CP47, and CP43 as well as TEF5 and PSB33 and their interactions using AlphaFold2 and 3 (Jumper et al., 2021; Abramson et al., 2024), which produced similar results. This analysis revealed one stable position of TEF5 and PSB33 in complex with the respective PSII core (Figure 12), where the Rieske-like domain points into the stroma, consistent with what has been determined experimentally (Fristedt et al., 2015). The main specific interactions are between TEF5/PSB33 and CP47 but also interactions with D1 (TEF5) and D2 (TEF5 and PSB33) are involved in stabilizing the complex (Figure 12). Specific to TEF5 is a predicted inter-protein β-sheet formed between TEF5 and the N-terminus of D2 (Figure 12A) that is not formed by PSB33. Nevertheless, PSB33 and TEF5 are predicted to interact equally strong with the PSII core and to share comparable interaction interfaces (Supplemental Figure S18). AlphaFold could not predict an interaction of TEF5 with the D1/D2 core where TEF5 exhibits the correct topology with the Rieske-like domain pointing to the stroma. Such an interaction would be expected from the co-migration of TEF5 with RCII (Figures 6B, 7). Possibly, such interactions take place with Cyt*b_559_* or PsbI not present in our models or TEF5/PSB33 interactions require conformational changes in RCII that are not predicted by AlphaFold.

**Figure 12.**
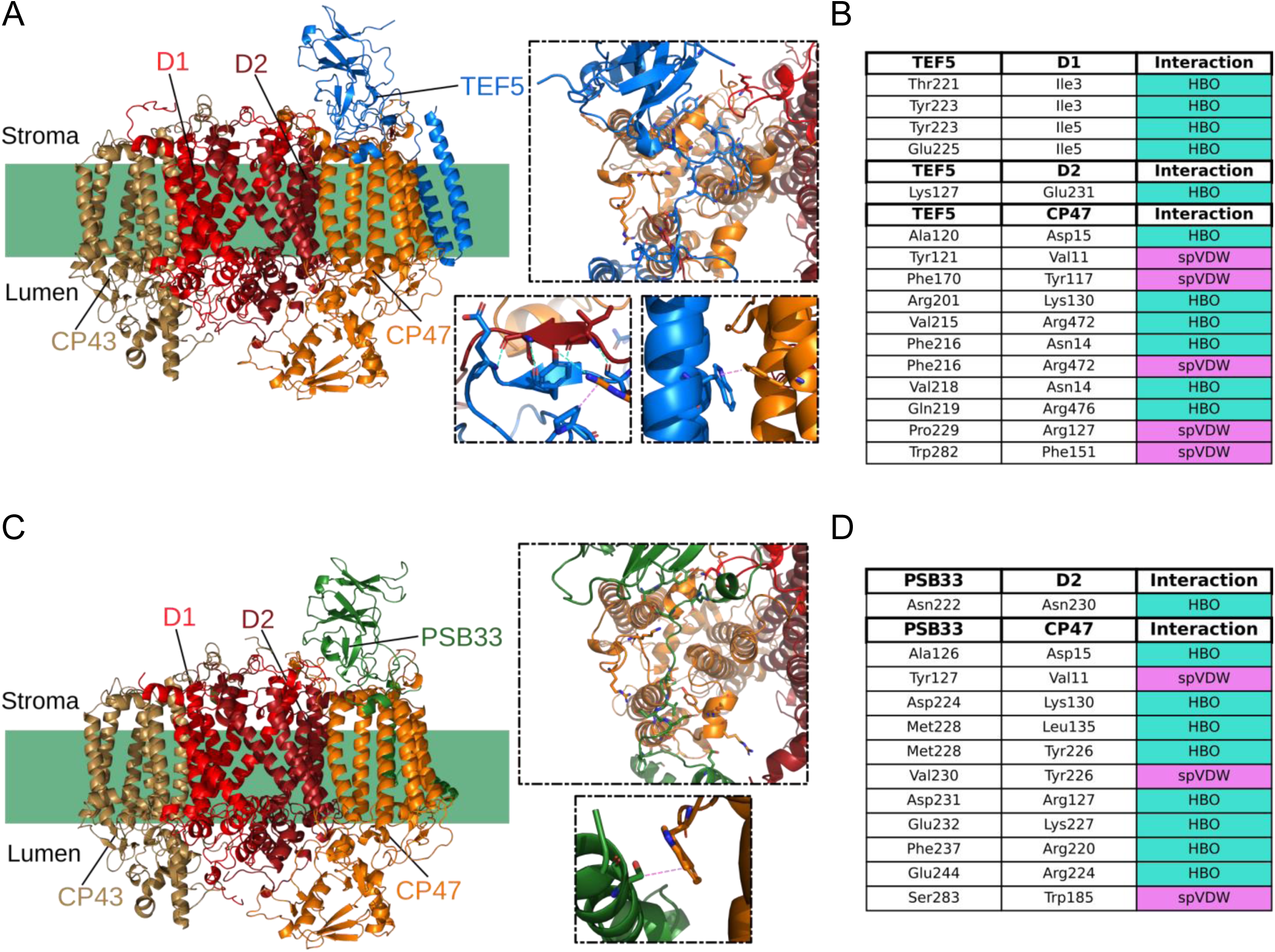
Predicted TEF5/PSB33-PSII core complexes. **(A)** Predicted structural model of TEF5 (blue) in complex with the PSII core (D1 – light red, D2 – dark red, CP47 – orange, CP43 – ochre) in Chlamydomonas. Highlighted are selected atomic interactions of the contact interfaces. **(B)** List of identified hydrogen bonds (HBO) and specific van-der-Waals interactions (spVDW) between TEF5 and PSII core subunits. **(C)** Predicted structural model of PSB33 (green) in complex with the PSII core (colored as in (A)) in Arabidopsis. Highlighted are selected atomic interactions of the contact interfaces. **(D)** List of identified hydrogen bonds (HBO) and specific van-der-Waals interactions (spVDW) between PSB33 and PSII core subunits.

The clearly reduced synthesis rates of CP47 and PsbH in the *psb28* and *tef5* mutants (Figures 3 and 10) suggests that a similar CES-like negative feedback control is at work in both. As was proposed above, this might be directly or indirectly triggered by RCII and CP43_mod_ accumulating in both, *psb28* and *tef5*. RCII and CP43_mod_ did not accumulate in the Chlamydomonas *lpa2* mutant, where no reduced translation rates were observed for any PSII core subunit (Spaniol et al., 2022).

## Materials and Methods

### Strains and culture conditions

*Chlamydomonas reinhardtii* wild-type CC-4533 and mutant strains LMJ.RY0402.193950 (*psb28*) and LMJ.RY0402.242855 (*tef5*) from the *Chlamydomonas* library project (Li et al., 2016) were obtained from the *Chlamydomonas* Resource Center. *psb28* and *tef5* mutants were used as recipient strains for transformation with plasmids pMBS687, pMBS703, and pMBS756 to generate complemented lines *psb28*-c2 and *psb28*-c6, and *tef5*-c15 and *tef5*-HA. Transformation was done via agitation with glass beads (*psb28* mutant) (Kindle, 1990) and electroporation (*tef5* mutant) (Shimogawara et al., 1998). Unless indicated otherwise, cultures were grown mixotrophically in TAP medium (Kropat et al., 2011) on a rotatory shaker at 25°C and ∼30 µmol photons m^-2^ s^-1^ provided by MASTER LEDtube HF 1200 mm UO 16W830 T8 and 16W840 T8 (Philips). For high-light exposure, cells were grown to a density of 2∼10^6^ cells mL^-1^, transferred to an open 1-L beaker, placed on an orbital shaker, and exposed to 1,200 to 1,800 µmol photons m^-2^ s^-1^ provided by CF Grow (CXB3590-X4). Cell densities were determined using a Z2 Coulter Counter (Beckman Coulter). For spot tests, cells were grown to a density of 3-5 x 10^6^ cells mL^-1^ and diluted in TAP medium such that 10 µl contained 10^4^, 10^3^ or 10^2^ cells. 10 µl of each dilution were spotted onto agar plates containing TAP medium or HSM medium and incubated in low light (30 µmol photons m^-2^ s^-1^) for 72 h, high light (600 µmol photons m^-2^ s^-1^) for 72 h, or in the dark for 96 h. HSM was prepared according to Sueoka (1960), but using the trace solutions from Kropat et al. (2011).

### Cloning of constructs for complementing the *psb28 and tef5* mutants

The *Chlamydomonas PSB28* coding sequence, including both introns, was amplified by PCR from *Chlamydomonas* genomic DNA in two fragments of 715 bp and 204 bp to remove an internal BsaI site using primers PSB28-1/2 and PSB28-3/4, respectively (Supplemental Table S1). The PCR products were cloned into the recipient plasmid pAGM1287 (Weber et al., 2011) by restriction with BbsI and ligation with T4-DNA ligase, resulting in the level 0 construct pMBS685. The *Synechocystis psb28-1* coding sequence, interrupted by the first *RBCS2* intron, was synthesized by BioCat (Heidelberg) with optimal *Chlamydomonas* codon usage and cloned into pAGM1287, yielding level 0 construct pMBS695. The *Chlamydomonas TEF5* coding sequence, interrupted by the first two *RBCS2* introns, was synthesized by BioCat (Heidelberg) and cloned into pAGM1287, giving level 0 construct pMBS701. The B3-B4 level 0 parts with the coding sequences were then complemented with level 0 parts (pCM) from the *Chlamydomonas* MoClo toolkit (Crozet et al., 2018; Niemeyer et al., 2021) to fill the respective positions in level 1 modules as follows: A1-B1 – pCM0-015 (*HSP70A-RBCS2* promoter + 5’ UTR), A1-B2 – pCM0-020 (*HSP70A-RBCS2* promoter + 5’ UTR), B2 – pMBS640 (CDJ1 chloroplast transit peptide); B5 – pCM0-100 (3xHA) or pCM0-101 (MultiStop); B6 – pCM0-119 (*RPL23* 3’UTR). The level 0 parts and destination vector pICH47742 (Weber et al., 2011) were directionally assembled into level 1 modules pMBS686 (PSB28-3xHA), pMBS696 (SynPsb28-1-3xHA), pMBS702 (TEF5-MultiStop), and pMBS755 (TEF5-3xHA) with BsaI and T4-DNA ligase. Level 1 modules were then combined with pCM1-01 (level 1 module with the *aadA* gene conferring resistance to spectinomycin), with plasmid pICH41744 containing the proper end-linker, and with destination vector pAGM4673 (Weber et al., 2011), digested with BbsI, and ligated to yield level 2 devices pMBS687 (PSB28-3xHA), pMBS697 (SynPsb28-3xHA), pMBS703 (TEF5-MultiStop), and pMBS756 (TEF5-3xHA). All MoClo constructs employed and generated are listed in Supplemental Table S3.

### Production of recombinant PSB28 in *E. coli*

The PSB28 coding region lacking the predicted chloroplast transit peptide (Figure 1A) was PCR-amplified from cDNA using oligonucleotides PSB28-Bam and PSB28-Hind (Supplemental Table S1). The resulting 438-bp PCR product was digested with BamHI and HindIII and cloned into the pETDuet vector (Novagen) (pMS1079), introducing an N-terminal 6xHis tag. Recombinant PSB28 was produced in *E. coli* ER2566 and purified by Ni-NTA affinity chromatography. Recombinant CGE1 was produced and purified as described previously (Willmund et al., 2007).

### Genotyping

3 x 10^7^ *Chlamydomonas* cells were centrifuged at 3500 *g* for 5 min. The pellet was resuspended in 250 μl water, followed by the addition of 250 μl 100 mM Tris-HCl pH 8, 10 mM EDTA, 4% SDS and incubation with proteinase K for 1 h at 55°C. Subsequently, 80 μl 5 M KCl and 70 μl CTAB/ NaCl (10% / 4%) were added, followed by agitation at 65°C for 10 min. DNA was extracted first with phenol / chloroform / isoamyl alcohol (25 : 24 : 1), then with chloroform / isoamyl alcohol (24:1). DNA was then precipitated with isopropanol and washed with 70% EtOH. The dried DNA pellet was dissolved in TE buffer (10 mM Tris-HCl pH 8, 1 mM EDTA) containing RNase. For PCR, genomic DNA, KAPA GC reaction buffer and KAPA Hifi HotStart Polymerase (Roche), 1 M betaine, 0.2 mM deoxynucleotide triphosphates, and 0.3 mM of the respective primers were mixed, incubated at 95 °C for 3 min and subjected to 35 cycles of 98°C for 20 sec, 63°C for 20 sec, and 72°C for 90 sec, followed by 75 sec at 72°C.

### qRT-PCR

RNA extraction and qRT-PCR analysis was done as described previously for the *lpa2* mutant (Spaniol et al., 2022) using the primers for *TEF5* and *CBLP2* as housekeeping control listed in Supplemental Table S1.

### SDS-PAGE and immunoblot analyses

Cells were harvested by centrifugation and frozen at -20°C. Frozen cell pellets were resuspended in sample buffer containing 62 mM Tris-HCl, pH 6.8, 2% (w/v) SDS and 10% (v/v) glycerol, boiled for 1 min at 95 °C, cooled on ice for 2 min, and centrifuged at 18,500 *g* and 25°C. Samples were diluted with sample buffer containing 50 mM DTT and 0.01% bromophenol blue to 1 µg protein µl^-1^ and subjected to SDS-PAGE and semi-dry western blotting. Antisera used were against D1 (Agrisera AS05 084), D2 (Agrisera AS06 146), CP43 (Agrisera AS11 1787), CP47 (Agrisera AS04 038), LHCBM9 (M. Schroda, unpublished data), PsaA (Agrisera AS06 172), PSAD (Agrisera AS09 461), PSAN (M. Schroda, unpublished data), Cyt *f* (Pierre and Popot, 1993), CGE1 (Schroda et al., 2001), CF1β (Lemaire and Wollman, 1989), RPL1 (Ries et al., 2017), and the HA-tag (Sigma-Aldrich H3663). Peptide antibodies against PSB28 and TEF5 were produced by Pineda (Berlin). Anti-rabbit-HRP (Sigma-Aldrich) was used as secondary antibody. Densitometric band quantifications after immunodetections were done with the FUSIONCapt software.

### Pulse-chase labeling

Cells in the exponential growth phase (2 x10^6^ cells mL^-1^) from a 100-mL culture were harvested by centrifugation, washed with minimum medium and resuspended in 1/20th volume of minimum medium. Cells were allowed to recover and to deplete their intracellular carbon pool for 1.5 hours under dim light (20 µE m^-2^ s^-1^) and strong aeration at 25°C. 10 µM cycloheximide and 10 µCi mL^-1^ Na-^14^C acetate (PerkinElmer: 56.6 mCi mM^-1^) were then added to the culture for the 7-min pulse. Cell samples, collected immediately after centrifugation at 4°C, were resuspended in ice-cold 0.1 M dithiothreitol and 0.1 M Na_2_CO_3_, frozen in liquid nitrogen, and kept at –80°C until analysis. For chase experiments, pulse-labelled cells were diluted in 35 ml of TAP medium containing 50 mM non-radioactive acetate and 250 μg ml−1 chloramphenicol at 25°C and further incubated in this medium for 20 and 60 min. Cells were then collected by centrifugation at 4°C and treated as above.

### BN-PAGE

BN-PAGE was performed with minor modifications according to (Jarvi et al., 2011). For the analysis of whole-cell proteins, 2 x 10^8^ cells (or 60 µg isolated thylakoids, see below) were centrifuged for 5 min at 4,400 *g*, 4°C, and resuspended in 750 μL of TMK buffer (10 mM Tris-HCl pH 6.8, 10 mM MgCl_2_, 20 mM KCl). After a further centrifugation step for 2 min at 2,150 *g*, 4°C, the pellet was resuspended in 350 µL ACA buffer (750 mM ɛ-aminocaproic acid, 50 mM bis-Tris/HCl pH 7.0, 0.5 mM EDTA), mixed with 4 μL of 25-fold protease inhibitor (Roche), and frozen at 80°C. The sample was then thawed on ice and sonicated for 30 sec (output: 25%, cycle: 70%), followed by a 5-min centrifugation at 300 *g* and 4°C. The protein concentration of the supernatant was determined according to (Bradford, 1976) and the sample was diluted with ACA buffer to 1.2 µg protein µL^-1^. For solubilization, 225 µL of the sample were mixed with 25 µL 10% β-DDM and incubated on ice for 20 min. After a centrifugation for 10 min at 18,500 *g* and 4°C, 15 μL loading buffer (250 mM ɛ-aminocaproic acid, 75% glycerol, 5% Coomassie Brilliant Blue 250 G) was added to the supernatant and samples were centrifuged several times at 18,500 *g* and 4°C until insoluble material was no longer present. Samples were then loaded on 4-15% BN acrylamide gels. Gels were either stained with Coomassie Brilliant Blue or the protein complexes were transferred to PVDF membranes. For the latter, the gel was incubated for 30 min in T2 buffer (25 mM Tris-HCl pH 10.4, 20% isopropanol) containing 0.1% SDS, then for a further 15 min in T2 buffer without SDS. The PVDF membrane (0.45 μm) was soaked in methanol for 15 sec and washed twice for 5 min with water. The membrane was then incubated in T2 buffer for 10 min. Proteins were transferred onto the membrane by semidry blotting using T1 buffer (25 mM Tris-HCl pH 9.8, 40 mM ɛ-aminocaproic acid, 20% isopropanol) containing 0.01% SDS.

For complexome profiling, thylakoids were isolated according to (Chua and Bennoun, 1975) with minor modifications. Briefly, 2 x 10^9^ cells were pelleted and washed with 25 mM HEPES-KOH, pH 7.5, 5 mM MgCl_2_ and 0.3 M sucrose, before resuspending in the same buffer supplemented with protease inhibitor (Roche). Cells were then lysed using a BioNebulizer (Glas-Col) with an operating N_2_ pressure of 1.5 bar. After centrifugation at 3,500 *g* for 10 min, the pellet was washed with 5 mM HEPES-KOH, pH 7.5, 1 mM EDTA and 0.3 M sucrose before resuspending in 5 mM HEPES-KOH, pH 7.5, 1 mM EDTA and 1.8 M sucrose. After placing 1.3 and 0.5 M sucrose layers in the same buffer on top and centrifugation at 100,000 *g* for 1 h, intact thylakoids, floating between the 1.3 M and 1.8 M layers, were collected, and diluted with 5 mM HEPES-KOH, pH 7.5 and 1 mM EDTA.

### In-gel digestion and mass spectrometry

Coomassie stained BN-PAGE gel pieces were destained by repeated cycles of washing with 40 mM NH₄HCO₃ for 5 min and incubating in 70% acetonitrile for 15 min, until they were colorless. They were then dehydrated completely by adding 100% acetonitrile for 5 min and dried under vacuum. Samples were then digested by covering the gel pieces in 10 ng/µl trypsin in 40 mM NH₄HCO₃ and incubating them over night at 37 °C, before first, hydrophilic peptides were extracted with 10% acetonitrile and 2% formic acid for 20 min and afterwards, all other tryptic peptides were extracted with 60% acetonitrile and 1% formic acid. Samples were combined and desalted according to (Rappsilber et al., 2007). Mass spectrometry was performed as described previously (Hammel et al., 2018; Spaniol et al., 2022).

### Evaluation of MS data

The analysis of MS runs was performed using MaxQuant version 1.6.0.16 (Cox and Mann, 2008). Library generation for peptide spectrum matching was based on *Chlamydomonas reinhardtii* genome release 5.5 (Merchant et al., 2007) including chloroplast and mitochondrial proteins. Oxidation of methionine and acetylation of the N-terminus were considered as peptide modifications. Maximal missed cleavages were set to 3 and peptide length to 6 amino acids, the maximal mass to 6000 Da. Thresholds for peptide spectrum matching and protein identification were set by a false discovery rate (FDR) of 0.01. The mass spectrometry proteomics data have been deposited to the ProteomeXchange Consortium via the PRIDE (Perez-Riverol et al., 2019) partner repository with the dataset identifier PXD023478. Total protein group intensities varied between samples. For sample normalization, the total ion intensity sum (TIS) of every protein and gel slice was calculated for each of the six samples (3x WT and 3x mutant). Sample normalization was performed by aligning protein group intensities of ATP synthase subunits ATPC, atpI, atpE, atpF, atpB, atpA, ATPD, and ATPG using the median of ratios method (Love et al., 2014). This resulted in a single correction factor for each sample. Subsequently, every intensity value was divided by its sample specific correction factor, to equalize all TISs. For further analysis, proteins identified by non-proteotypic peptides were discarded. Protein identifiers were annotated with MapMan ontology terms, Gene Ontology (GO) terms, and proposed subcellular localization (https://doi.org/10.5281/zenodo.6340413). A Welch test was performed for each protein by considering the sums of all 36 normalized slice intensities for each sample and testing three WT sums against three mutant sums. The distance of the average migration profiles for every protein was calculated as the Euclidean distance between WT and mutant. To adjust for amplitude-introduced bias, each distance was divided by the maximal average intensity of WT or mutant, respectively. Data normalization and analysis were performed using FSharp.Stats (https://doi.org/10.5281/zenodo.6337056). The migration profiles were visualized using Plotly.NET (Schneider et al., 2022).

### Immunoprecipitation

200 ml of culture was grown in HAP medium (TAP in which Tris was replaced by 20 mM HEPES-KOH pH 7.0) and supplied for 10 min with formaldehyde (0.37% final concentration) for *in-vivo* crosslinking. 100 mM Tris-HCl pH 8.0 was added to the culture for quenching before cells were collected by centrifugation for 5 min at 2500 *g* and 4°C. The cell pellet was resuspended in 1.5 mL TE buffer and frozen at -20°C. After thawing at 23°C, 20 μL PMSF was added and samples were frozen in liquid nitrogen. After two more cycles of thawing and freezing, 50 μL were taken to determine the protein concentration and samples were centrifuged for 30 min at 18,000 *g* and 4 °C. 40 mM Tris-HCl pH 8, 150 mM NaCl, 1 mM MgCl_2_, 10 mM KCl, and 0.1% α-DDM were then added to the supernatant. The pellet was resuspended in TNMK buffer (50 mM Tris-HCl pH 8, 150 mM NaCl, 1 mM MgCl_2_, 10 mM KCl) containing 1% α-DDM. Samples were then mildly sonicated and centrifuged at 14,000 *g* and 4°C for 10 min after a 5-min incubation on ice. The supernatants were added to 20 μL HA-coupled magnetic beads (Pierce) and the samples were incubated for 1.5 h at 4 °C. After three washing steps with TNMK buffer containing 0.05% Tween and three washing steps with TNMK buffer, 100 μL of sample buffer (90 mM Tris-HCl, 20% glycerol, 2% SDS) were added and the samples were boiled for 1 min. The eluate was removed from the magnetic beads, mixed with 50 mM DTT and boiled for an additional 10 min. The eluates were then analyzed by SDS-PAGE and immunodetection or by mass spectrometry.

### Chlorophyll fluorescence measurements

Chlorophyll fluorescence was measured using a pulse amplitude-modulated Mini-PAM fluorometer (Mini-PAM, H. Walz, Effeltrich, Germany) essentially according to the manufacturer’s protocol after 3 min of dark adaptation (1 s saturating pulse of 6,000 μmol photons m^-2^ s^-1^, gain = 4).

### Chlorophyll precursors

The tetrapyrrole biosynthesis intermediates and end-products were analyzed by High Pressure Liquid Chromatography (HPLC), essentially as described previously (Brzezowski et al., 2014), on cultures grown in dark or in low light (30 μmol photons m^−2^ s^−1^). In short, samples containing 1.2 × 10^8^ cells were centrifuged at 3000 *g* for 5 min at 4 °C and the pellets were snap-frozen in liquid N_2_. Protoporphyrin IX, Mg-protoporphyrin IX, Mg-protoporphyrin IX monomethylester, protochlorophyllide, chlorophyllide, Chl a and b, and pheophorbide were extracted in 500 µL cold (-20 °C) acetone/0.1M NH_4_OH (9/1, v/v) with sonication, followed by a three-step cycle of resuspension and centrifugation using the same solution. Heme extraction was performed on the remaining pellet using 100 µL acetone/HCl/DMSO (10/0.5/2, v/v/v) in the same three-step protocol. HPLC analyses were performed essentially as described in (Czarnecki et al., 2011). Values were normalized to pmol/10^6^ cells.

### P700 decay kinetics

Measurements of P700+ reduction kinetics were conducted using a Dual-PAM-100 instrument from Heinz Walz (Effeltrich, Germany), with chlorophyll concentrations set to 5 µg/mL. To fully oxidize P700, a 50-millisecond multiple-turnover light pulse at an intensity of 10,000 µmol photons m^−2^ s^−1^ was administered following a 2-min dark incubation period. For WT, *psb28*, and complemented lines, the reduction kinetics of P700+ were monitored without any additions and in the presence of 100 µM DCMU. The average results from four separate experiments were fitted with single exponential functions and analyzed as described previously (Bernát et al., 2009). The presented values were calculated using one-way ANOVA analysis with the GraphPad Prism software.

### Light and transmission electron microscopy

Light microscopy images were taken with an Olympus BX53F microscope with 100x magnification. For transmission electron microscopy, cells were collected and washed in 100 mM sodium cacodylate at pH 7.2. Afterwards, cells were fixed in 100 mM sodium cacodylate containing 2.5% glutaraldehyde and 4% formaldehyde at pH 7.2 at room temperature. The buffer was exchanged after 20 min, 60 min and 120 min. All other steps were done as described previously (Nordhues et al., 2012). Samples were analyzed with a JEM-2100 (JEOL) transmission electron microscope (operated at 80 kV). Micrographs were taken using a 4,080-3 4,080-pixel CCD camera (UltraScan 4000; Gatan) and the Gatan DigitalMicrograph software (version 1.70.16).

### Immunofluorescence microscopy

Formaldehyde was added to a final concentration of 4% to 1 ml *Chlamydomonas* cells grown to log phase, followed by an incubation at 4 °C for 1 h. 10 μl of 0.1% poly-L-lysine were applied to a microscopy slide and 40 μl of fixed cells were added. The slide was then placed into ice-cold methanol for 6 min. Subsequently, the slide was washed five times with PBS. For permeabilization, cells were incubated in PBS containing 2% Triton at 25°C. After five more washing steps with PBS containing 5 mM MgCl_2_, the slide was incubated in PBS containing 1% BSA and the primary antibody was added (rabbit anti-D1, Agrisera AS05 084, 1:10,000; mouse anti-HA, Pineda, 1:3,000), followed by an incubation overnight. After five washes with PBS containing 1% BSA, the secondary antibody (fluorescine-isothiocyanate-labeled goat anti-rabbit (Sigma-Aldrich); Alexa Fluor 488 goat anti-mouse (Thermo Fisher Scientific), 1:500) was added followed by an incubation for 1.5 h. Five last washes with PBS followed before microscopy images were taken. To this end, a Zeiss LSM880 AxioObserver confocal laser scanning microscope equipped with a Zeiss C-Apochromat 40×/1,2 W AutoCorr M27 water-immersion objective was used. Fluorescent signals of FITC (excitation/emission 488 nm/493–553 nm) and Alexa Fluor 546 (excitation/emission 543 nm/553–669 nm) were processed using the Zeiss software ZEN 2.3 or Fiji software. For double labeling, images were acquired using sequential scan mode to avoid channel crosstalk.

### Structural Modeling

The complexes formed by TEF5 and PSB33 with the respective PSII cores were predicted by AlphaFold2 (Jumper et al., 2021) and AlphaFold3 (Abramson et al., 2024). We used a local AlphaFold2 installation through ColabFold (Mirdita et al. 2022) to predict five different complexes. Models were ranked based on the ColabFold scoring tools IDDT (Mariani et al., 2013) and TM (Zhang and Skolnick, 2004). The predicted structures were subsequently minimized utilizing the AMBER force field optimization (Weiner et al., 1984). Analogously, the complexes were predicted using the AlphaFold3web server. As AlphaFold2 and AlphaFold3 gave comparable results, only the best ranked AlphaFold3 structures were used for further interaction evaluation. The contact network of the assembly factors TEF5 and PSB33 with RCII were analyzed with MAXIMOBY/MOBY (CHEOPS, Germany) and PyContact (Scheurer et al., 2018).

### Sequence alignments, motif search and pairwise structure comparisons

Putative chloroplast transit peptides of PSB28 and TEF5 homologs were predicted with TargetP (Almagro Armenteros et al., 2019) and putative transmembrane helices with DeepTMHMM (Hallgren et al., 2022). Sequence motifs were searched by InterProScan (Jones et al., 2014). Pairwise structural comparisons were done with the Analyze tool in RCSB (https://www.rcsb.org/) (Berman et al., 2000) and displayed with Mol* (Sehnal et al., 2021). Sequence alignments were done with CLUSTALW (https://www.genome.jp/tools-bin/clustalw) and displayed with GeneDoc.

## Supplemental Files

**Supplemental Figure S1.** Analysis of the CIB1 integration site in the *PSB28* gene by PCR and testing of the PSB28 peptide antibody.

**Supplemental Figure S2.** Screening for complemented *psb28* transformants.

**Supplemental Figure S3.** Analysis of chlorophyll precursor contents in *psb28* mutant and WT.

**Supplemental Figure S4.** Analysis of PSII complex assembly, subunit accumulation, and functionality in dark-grown cells.

**Supplemental Figure S5.** Analysis of oligomerization capacity of recombinant PSB28 and quantification of cellular PSB28 abundance.

**Supplemental Figure S6.** Monitoring kinetics of PSII repair after photoinhibition in the *psb28* mutant.

**Supplemental Figure S7.** Monitoring kinetics of PSII re-synthesis in the *psb28* mutant after sulfur starvation.

**Supplemental Figure S8.** Construct for the expression of *Synechocystis* Psb28-1 and analysis of transformants in the *psb28* mutant background.

**Supplemental Figure S9.** BN-PAGE for complexome profiling.

**Supplemental Figure S10.** Comparison of BN-PAGE migration profiles of PSII core subunits and of putative novel PSII-associated protein PBA1.

**Supplemental Figure S11.** Comparison of BN-PAGE migration profiles of thylakoid membrane protease FTSH1/2 and kinase STL1.

**Supplemental Figure S12.** Comparison of BN-PAGE migration profiles of PSII core subunits, of known PSII auxiliary factors, and of putative novel auxiliary factors.

**Supplemental Figure S13.** Analysis of the CIB1 integration site in the *TEF5* gene by PCR and of TEF5 protein in the *tef5* mutant and in complemented lines.

**Supplemental Figure S14.** Screening for complemented *tef5* transformants.

**Supplemental Figure S15.** Monitoring kinetics of PSII repair after photoinhibition.

**Supplemental Figure S16.** Monitoring kinetics of PSII re-synthesis in the *tef5* mutant after sulfur starvation.

**Supplemental Figure S17.** Alignment of N-terminal regions of green algal LHL4 and PSBS proteins with cyanobacterial HliA-D.

**Supplemental Figure S18.** Predicted contact interfaces of TEF5/PSB33 with PSII cores in Chlamydomonas and Arabidopsis.

**Supplemental Table S1.** Primers used for genotyping, cloning, and RT-PCR.

**Supplemental Table S2.** Proteins involved in PSII assembly, repair, or complex dynamics that have clear homologs in Chlamydomonas and are present with three replicates each for WT and *psb28* mutant in the complexome profiling dataset.

**Supplemental Table S3.** MoClo constructs employed and generated.

**Supplemental Data Set S1.** LC-MS/MS analysis of PSB28 immunoprecipitates.

**Supplemental Data Set S2.** Interactive complexome profiling dataset.

**Supplemental Data Set S3.** Heat maps of all proteins found in the complexome profiling dataset in the WT and the *psb28* mutant.

**Supplemental Data Set S4.** LC-MS/MS analysis of TEF5 immunoprecipitates.

## Acknowledgements

This work was supported by the Deutsche Forschungsgemeinschaft [FOR2092, SFB/TRR175, project C02] and the Profilbereich BioComp. We would like to thank Karin Gries for technical assistance.

## Author Contributions

J.L. and K.K. performed all experiments supported by B.S. and L.S. B.V. evaluated the complexome profiling data supervised by T.M. F.S. generated all mass spectrometry data. M.M. performed the pulse-chase experiments supervised by Y.C. and F.-A.W. S.G. recorded the electron microscopy images and D.S. recorded the confocal microscopy images. P.B. measured chlorophyll, precursors and breakdown products. J.Z. analyzed PSI kinetics supervised by M.N. T.F. modeled the PSII-TEF5/PSB33 structures supervised by T.R. M.S. conceived and supervised the project and wrote the article with contributions from all authors.

